# Synaptojanin 2 is required for neuronal mRNA localization, mitochondrial structure and function, and synaptic metabolism

**DOI:** 10.64898/2026.11.24.768930

**Authors:** Whitney S. Gibbs, Sindhuja Gowrisankaran, Thibaud Martial, Gagandeep Kaur, Sarah DeVries, Thomas L. Schwarz

## Abstract

Neurons rely on local mRNA transport to support the metabolic and structural demands of their distal compartments. Synaptojanin 2a (Synj2a), a mitochondria-associated phosphoinositide phosphatase, binds numerous mitochondrial mRNAs via an RNA recognition motif (RRM) and delivers them to axons and dendrites. Among these, PINK1 mRNA transport is particularly critical for mitochondrial quality control. Here we show that this RNA transport function, independent of Synj2’s catalytic activity, is required for PINK1-dependent mitochondrial quality control, synaptic bioenergetics, and neuronal resilience. We made mice and human iPSC-derived neurons carrying RRM point mutations that disrupt RNA binding while preserving both phosphatase domains. The mutations also reduced endogenous Synj2 protein levels, indicating that RNA binding contributes to Synj2 stability. Rescue experiments with the RRM targeted to the outer mitochondrial membrane were therefore necessary to isolate which effects are due to RNA-binding. Neurons with the mutated RRM failed to localize PINK1 mRNA to neurites. Consequently, depolarizing mitochondria in these neurites failed to stabilize PINK1 protein or cause phosphoubiquitin to accumulate on mitochondria. Re-expressing the RRM rescued these defects, confirming that RNA-binding rather than phosphatase activity drives the phenotype. Mutant mice showed abnormal mitochondrial cristae ultrastructure and reduced respiratory capacity in synaptosomes. Metabolic profiling revealed extensive reprogramming, most notably increased glycolytic flux, elevated branched-chain amino acids and reduced cholesterol. Despite compensatory reprogramming, mutant neurons showed heightened vulnerability to mitochondrial and excitotoxic stress. This vulnerability was partially rescued by the mitochondrially-targeted RRM. These findings establish Synj2-mediated mRNA transport as a significant mechanism that supports mitochondrial homeostasis.

**Significance Statement:** Brain cells need to keep mitochondria, the cell’s energy-producing organelles, functioning properly even at sites far removed from the cell body. This can require local delivery of the genetic instructions for building and maintaining mitochondria. We show that a protein called synaptojanin 2 transports a set of mitochondria-destined mRNAs, including the gene PINK1, out to distant mitochondria so that damaged ones can be identified and cleared. Mutations in PINK1 cause Parkinson’s disease. When we disrupted this RNA transport function in mice and human neurons by mutating the protein synaptojanin 2, PINK1 could no longer reach and protect distant mitochondria. Neuronal mitochondria had abnormal structures, and the neurons had disrupted energy metabolism, cholesterol loss, and increased vulnerability to stresses. These findings reveal mRNA transport by synaptojanin 2 as an important mechanism for maintaining mitochondrial health in neurons, with implications for understanding Parkinson’s disease and other neurodegenerative conditions.

## INTRODUCTION

The extraordinary architecture of neurons requires cell biological specializations of pre- and post-synaptic compartments that may be far from the nucleus and cell body. These specializations include forms of membrane trafficking, energetic and metabolic requirements, and local protein translation to supplement transport from the cell body. To achieve these ends, synapses have a heightened need for many of the proteins that mediate these processes and, in some cases, specialized isoforms of proteins. The phosphotidylinositide phosphatases of the Synaptojanin family of proteins fall in this category. Synaptojanin 1 is enriched at presynaptic terminals to facilitate endocytosis of synaptic vesicle membranes(1, 2). Synaptojanin 2 (Synj2) is less well studied though it shares many structural features with Synaptojanin 1. One set of isoforms (Synj2b) are concentrated at synapses (3). Another isoform, Synj2a, is enriched in neurons over non-neuronal cells and targeted to mitochondria (4).

Both Synaptojanin 1 and Synj2 contain two well-studied, distinct catalytic domains: a Sac1 domain that hydrolyzes phosphatidylinositol 3-phosphate and 4-phosphate, and a 5-phosphatase domain that acts on phosphatidylinositol 4,5-bisphosphate. All Synaptojanin isoforms also harbor an RNA-recognition motif (RRM) whose physiological significance has received less attention. Recently, we reported that the mitochondria-associated Synj2a isoform can bind to multiple RNA species via its RRM and these include the transcript for PINK1 (PTEN-Induced Putative Kinase 1) and ATP51β (5). The set of mRNA species that coprecipitated with Synj2a upon its expression in HEK293T cells was enriched for mRNA encoding mitochondrial proteins (5) while other transcripts, such as that for β actin were not. Synj2a, via its RRM, could localize PINK1 mRNA to mitochondria and thereby allow the mRNA to be transported on mitochondria into axons and dendrites. The association of Synj2 with mRNA suggests a role in regulating translation of synaptic and mitochondrial proteins, including PINK1, in addition to its known phosphatase activities.

PINK1 serves as a critical sentinel of mitochondrial health; it accumulates on damaged mitochondria to initiate quality control responses including mitophagy(6). Loss of PINK1 function causes early-onset Parkinson’s disease, underscoring the importance of PINK1 regulation for neuronal survival(7). Because of PINK1’s role in mitochondrial quality, it is important to understand the mechanisms governing PINK1 mRNA localization and local translation at synaptic terminals. Given the short half-life of PINK1 protein(5, 8), the constraints of long-distance axonal transport, and the need for rapid responses to mitochondrial dysfunction, local PINK1 synthesis may be essential for effective quality control in distal neuronal compartments. In light of the multiple mRNA species for mitochondrial proteins that interact with Synj2, Synj2 is likely to have a major role in mitochondrial quality control and synaptic bioenergetics beyond PINK1 transport.

To determine the contribution of SYNJ2’s RNA-binding function to mitochondrial regulation, we generated mice and iPSC-derived human neurons harboring targeted mutations in the Synj2 RRM. These mutations were designed to preserve phosphatase activity while selectively disrupting RNA binding. Mutation of the RRM, however, led to a sharp decrease in Synj2 protein suggesting a requirement of RNA-binding for protein stability. We used these mice and iPSC-derived neurons to investigate the role of Synj2 in key aspects of mitochondrial homeostasis and neuronal physiology. To distinguish between phenotypes arising specifically from loss of RNA binding from those due to secondary loss of Synj2 protein, we employed complementation studies using RRM targeted to the outer mitochondrial membrane (OMM). Our findings reveal the importance of Synj2 for mitochondrial quality control, synaptic metabolism, synaptic energetics, and neuronal survival.

## RESULTS

### Generation and characterization of a mouse with a mutated SYNJ2 RRM

Substituting 3 alanines for Valine 909, Glutamine 951 and Leucine 953 within the RRM of rat Synj2a (Syn2^AAA^) had previously been shown to prevent the RNA-dependent shift in Synj2 motility after UV crosslinking and the ability of overexpressed RRM domains to target PINK1 transcripts to mitochondria (5). The RRM of Synj2 comes after the two phosphatase domains and before the alternatively spliced C-terminal region (Figure 1A). To determine whether mutations in the RRM also disrupt phosphatase function, we compared in HEK293T cells the catalytic activity of overexpressed Synj2^AAA^, WT, and Synj2a with mutations known to disrupt each phosphatase domain. All the proteins were expressed at comparable levels. As expected, mutation of the SAC domain (D388N) prevented hydrolysis of PI(3)P and PI(4)P and mutation of the 5-phosphatase domain (N538K) prevented hydrolysis of PI(4,5)P_2_. The VQL/AAA substitution in the RRM, however, retained phosphatase activity against each substrate equivalent to WT (Figure 1B,C), Thus, the RRM mutations disrupt RNA binding while preserving the activity of both catalytic domains.

**Figure 1.**
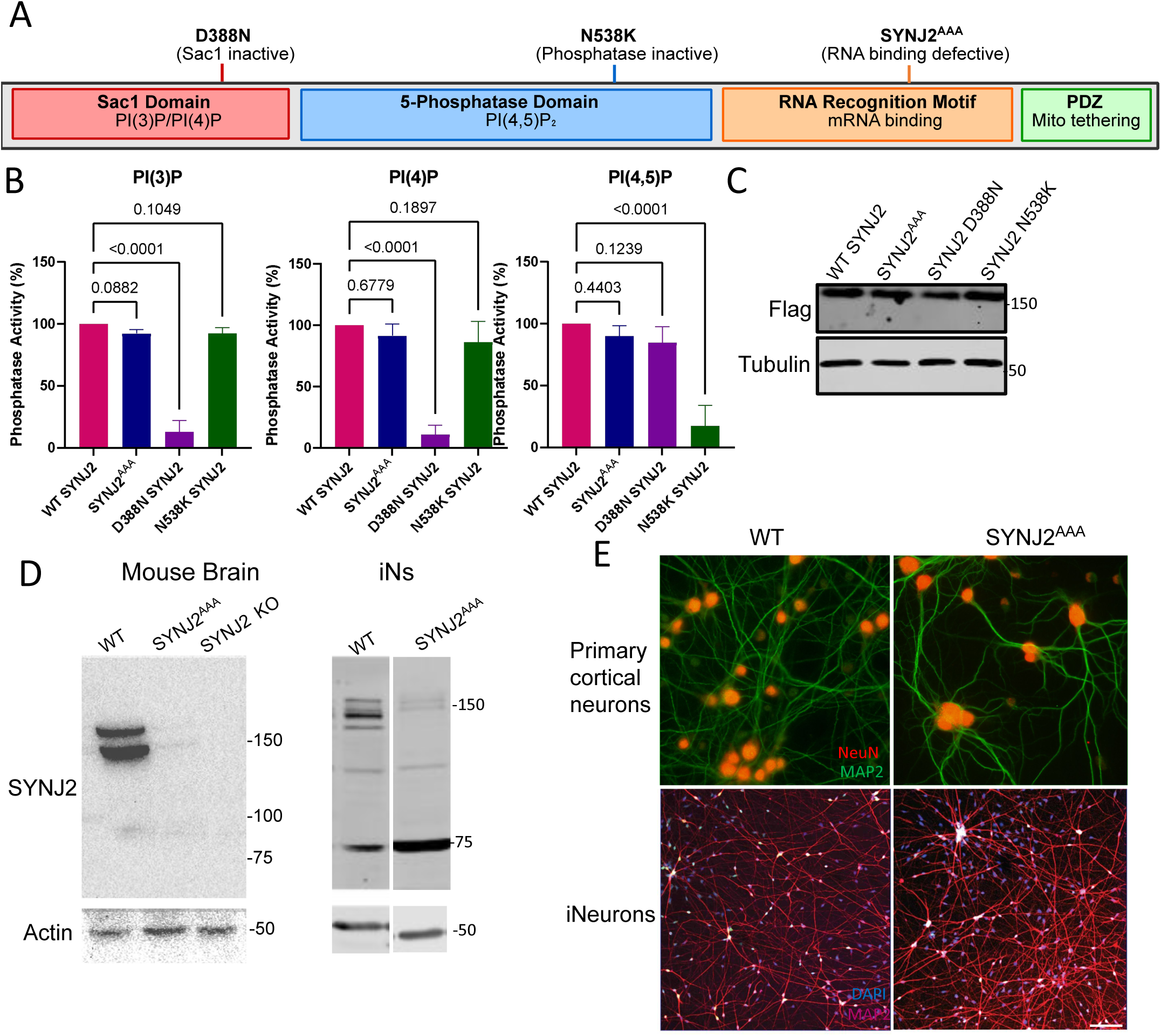
Generation and characterization of Synj2 RRM-domain mutants. **(A)** Schematic diagram illustrating the functional domains of synaptojanin 2 (Synj2), including the location of the RNA recognition motif (RRM) domain targeted for mutagenesis. **(B)** Phosphatase activity assay in HEK293T cells of wild-type (WT) Synj2 compared to Synj2^AAA^. SYNJ2 phosphatase-dead mutants, Synj2 D388N and SYNJ2 N538K serve as negative controls. Activity is expressed as percentage relative to WT based on hydrolysis of phosphoinositide substrates PI(3)P, PI(4)P, and PI(4,5)P₂. Statistical analysis by one way ANOVA with Tukey’s HSD multiple comparison test. n = 12. **(C)** Immunoblot of SYNJ2 protein in WT, Synj2^AAA,^ SYNJ2 D388N and Synj2 N538K expressed in HEK cells as in (B). **(D)** Immunoblot of endogenous Synj2 in whole mouse brain and iNeurons lysate from WT and SYNJ2^AAA^ samples probed with Synj2 antibody raised against the C terminal proline-rich domain of SYNJ2. Mouse brain immunoblot contains SYNJ2 KO control. The upper species (∼140-160kDa), which include the full-length Synj2 and its mitochondrial-localized isoform, are markedly reduced in Synj2^AAA^ relative to WT. In contrast, the ∼75-100kDa species are unchanged suggesting they correspond non-specific cross-reactive proteins. Actin serves as a loading control. **(E)** Upper panels show immunofluorescence microscopy of primary cortical neurons isolated from WT and Synj2^AAA^ mice and stained with NeuN (red, neuronal marker) and MAP2 (green, dendritic marker). Bottom panels show WT and Synj2^AAA^ iPSC-derived neurons stained with DAPI (blue, nuclei) and MAP2 (red, dendrites).

We therefore used CRISPR/Cas9 to make a Synj2^AAA^ mouse with mutations at the homologous amino acids in the RRM: V892A, Q934A, and L936A mutations. Sanger sequencing of isolated genomic DNA confirmed successful and precise introduction of all three mutations (Supplement Figure 1A). We also generated corresponding human iPSC lines with homozygous mutations in the equivalent amino acid residues (V892, Q934 and L936) and confirmed their sequences by Sanger sequencing (Supplemental Figure 1B). However, Western blot analysis of both mouse brain tissue and iPSC-derived neurons revealed an unexpected finding: the Synj2 RRM mutations caused near complete loss of endogenous Synj2 protein (Figure 1D) compared to WT controls, while RNA levels for total Synj2 and the mitochondrial-specific isoform (Synj2a) were unchanged (Supplement Figure 1C). The upper species (∼140-160kDa), which include the full-length Synj2 and its mitochondrial-localized isoform, are markedly reduced in SYNJ2^AAA^ relative to WT. This suggests that at endogenous expression levels, the RRM and its RNA-binding property play a critical role in Synj2 protein stability. This unexpected reduction in protein levels complicated the interpretation of the resulting phenotypes we subsequently uncovered: were they directly due to interactions with RNA or reflective of a general Synj2 loss-of-function, including loss of phosphatase activities? In experiments with cultured neurons, however, it was possible to circumvent the problem by asking if a phenotype could be rescued by expression of just the Synj2 RRM targeted to mitochondria. Despite the severe reduction in Synj2 protein levels, the Synj2*^AAA^*mice were homozygous viable and the mutations in Synj2 human iPSCs did not prevent their proliferation or neuronal differentiation and morphological development in culture, as demonstrated by normal NeuN and MAP2 staining patterns (Figure 1E).

### SYNJ2 RRM mutations disrupt PINK1 mRNA localization and mitochondrial quality control

To determine how mutation of the RRM altered the distribution of a transcript, we examined the subcellular localization of PINK1 mRNA, which we had previously shown to bind to Synj2(5). By combining RNAscope and immunocytochemistry for ATP5a to mark mitochondria, we analyzed cultured WT and mouse *Synj2*^AAA^ cortical neurons (Figure 2). WT neurons showed robust PINK1 mRNA localization to neurites and clear overlap with mitochondria. In contrast, *Synj2*^AAA^ neurons contained significantly less PINK1 mRNA in neurites and what RNAscope signal remained was less colocalized with mitochondria (Figure 2A, B), consistent with a defect in mRNA transport. Synj2 is normally localized to mitochondria by virtue of its interaction with Synaptojanin2 Binding Protein (Synj2BP, also called OMP25), an intrinsic protein of the mitochondrial outer membrane (4). To selectively restore the RNA-binding property of Synj2 to the mitochondrial surface we used a construct (mito-RRM) in which the RRM of mouse Synj2 was directly coupled to the transmembrane domain of OMP25.

**Figure 2.**
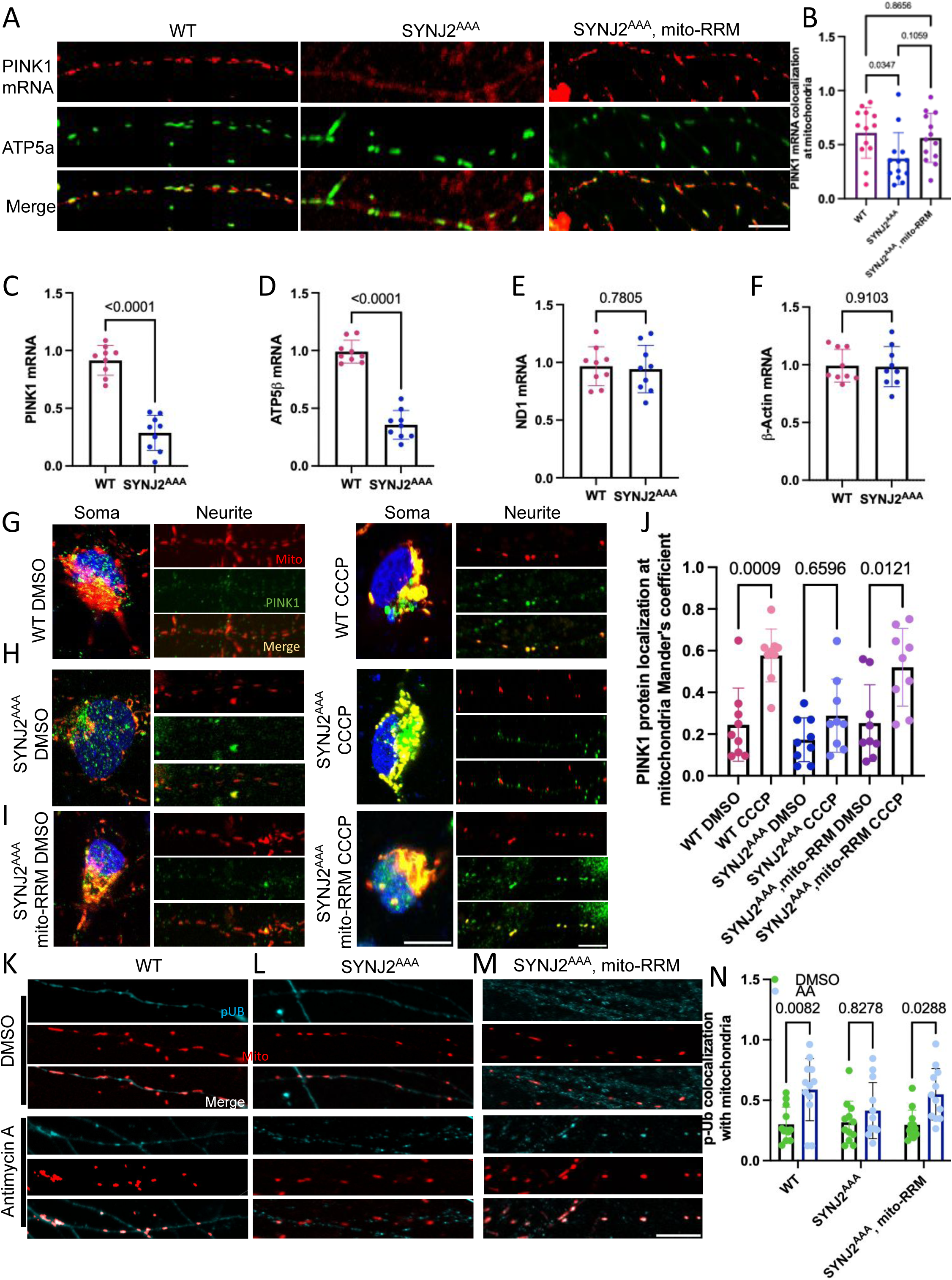
Synj2 RRM mutations disrupt PINK1 mRNA localization and mitochondrial quality control. **(A)** RNAscope in situ hybridization for PINK1 mRNA (red) followed by immunocytochemistry for ATP5A (mitochondrial marker, green) in primary cortical mouse neurons from WT, SYNJ2^AAA^, and SYNJ2^AAA^ neurons with mito-RRM overexpression. (**B)** Quantification of PINK1/ATP5A colocalization in neurites (Manders coefficient) for WT, Synj2^AAA^ and Synj2^AAA^, mito-RRM neurons from RNAscope experiment as in (A). Statistical analysis by one-way ANOVA with Tukey’s HSD multiple comparison test. n = 12-15. (**C-F)** Quantitative RT-PCR analysis of transcript levels in synaptosomes isolated from WT and Synj2^AAA^ brains for PINK1 **(C)**, ATP5A **(D)**,ND1 (mitochondrial-encoded NADH dehydrogenase subunit 1) **(E)**, and β-actin **(F).** Actin was used as a loading control n=9. Data expressed as fold-change relative to WT. **n** = 9 error bars represent SD. (**G-I)** Immunocytochemistry against PINK1 (green) and ATP5A (mitochondrial marker, red) in primary cortical WT **(G)**, Synj2^AAA^ **(H)**, and Synj2^AAA^ neurons expressing mito-RRM **(I)**, treated with DMSO or 10uM CCCP for 3 hrs to stabilize PINK1 protein. (**J)** Quantification of PINK1/ATP5A colocalization in neurites (Manders coefficient) for WT, Synj2^AAA^, and Synj2^AAA^, mito-RRM neurons under DMSO and CCCP conditions. Statistical analysis by one way ANOVA with Tukey’s HSD multiple comparison test. n = 12-15.(**K-M)** Immunocytochemistry against phospho-ubiquitin(pUb,cyan) and ATP5A(red) in WT **(K)**, Synj2^AAA^ **(L)**, Synj^AAA^ neurons overexpressing mito-RRM **(M)**, treated with DMSO or 40uM antimycin A for 2 hrs to trigger PINK1-dependent accumulation of phospho-ubiquitin on mitochondria. (**N)** Colocalization of phospho-Ubiquitin and mitochondrial ATP5A as in **(K-M)** quantified using Manders overlap coefficient. Statistical analysis by one-way ANOVA with Tukey’s HSD multiple comparison test. **n** = 12-15.

Overexpression of mito-RRM successfully rescued PINK1 mRNA localization to mitochondria in neurites although the construct lacks both Sac1 and 5-phosphatase domains (Figure 2A, B). Global PINK1 mRNA levels remained unchanged in *Synj2*^AAA^ neurons, confirming that the observed phenotype reflects impaired mRNA transport to neurites rather than transcriptional downregulation or instability.

We next examined whether this defect in mRNA localization extended to synaptic compartments in adult tissue by analyzing synaptosomes isolated from 6-month-old WT and *Synj2*^AAA^ mouse brains. Quantitative PCR analysis of *Synj2*^AAA^ synaptosomes revealed significantly reduced PINK1 and ATP51β mRNA levels, two transcripts known to interact with Synj2, compared to WT controls (Figure 2C, D). The mRNA for ND1, a mitochondrially-encoded gene, and β actin (Figure 2E, F), which does not bind to Synj2, showed no difference between genotypes, as expected for a specific transport defect rather than a global loss of mitochondria or mRNA transport.

The reduction in PINK1 mRNA localization was predicted to produce corresponding defects in PINK1 protein distribution and function. Under normal conditions, PINK1 protein is constitutively imported into healthy mitochondria, where it is rapidly cleaved by mitochondrial proteases that trigger its degradation. Cells with healthy mitochondria thereby maintain low steady-state levels of PINK1(6, 9). However, mitochondrial depolarization blocks PINK1 import, leading to its accumulation and stabilization on the outer mitochondrial membrane where it becomes catalytically active and initiates mitophagy. To assess this stress-response pathway, we turned to iPSC-derived cortical-like iNeurons to take advantage of the availability of antibodies to human PINK1. CCCP, a mitochondrial uncoupler that depolarizes mitochondria, can be used to trigger PINK1 stabilization(6). As expected, under basal conditions (DMSO treatment), immunolocalization of PINK1 and ATP5a (as a mitochondrial marker) in Synj2^AAA^ iNeurons or their isogenic parental line (WT) showed minimal PINK1 protein in either their soma or their neurites (Figure 2G, J). Upon treatment with 20 μM CCCP for 3 hours, WT iNeurons exhibited robust PINK1 protein stabilization on mitochondria in both their soma and neurites (Figure 2G, J). In contrast, in SYNJ2^AAA^ iNeurons, only the somatic mitochondria accumulated PINK1 protein; little or none was detected on mitochondria in neurites (Figure 2H, J). The selective loss of PINK1 accumulation in neurites is consistent with the accumulation depending on local synthesis of PINK1 which in turn is dependent on Synj2-dependent mRNA transport. Overexpression of the Synj2 mito-RRM construct in Synj2^AAA^ neurons restored PINK1 protein expression on distal mitochondria following CCCP treatment (Figure 2I,J), demonstrating that the RRM, and not the rest of the Synj2 protein, is necessary.

To assess whether PINK1 mRNA transport was essential for a functional PINK1 pathway in distal neurites of mouse primary neurons, we examined levels of mitochondrial phosphoubiquitin, a downstream readout of the mitophagy pathway activated by the kinase activity of PINK1. After inhibition of the electron transport chain with 40 μM Antimycin A, we imaged and quantified phosphoubiquitin accumulation on mitochondria by immunostaining. Phosphoubiquitin accumulated robustly in neurites of WT (Figure 2K, N) but not *Synj2*^AA*A*^ (Figure 2L, N) primary neurons. Again, rescue with the mitochondrially targeted mito-RRM restored phosphoubiquitin accumulation on mitochondria in *Synj2*^AAA^ neurons (Figure 2M, N), directly linking RRM-dependent mRNA transport and localization to functional mitochondrial quality control. Collectively, these findings reveal that axonal PINK1-dependent mitophagy is highly dependent on local translation of Synj2 and demonstrates the functional importance of the transport of its mRNA.

### SYNJ2 RRM mutations impair mitochondrial respiratory function and ultrastructure

Having established the significance of the RRM for mRNA transport *in vitro*, we next examined consequences of the Synj2^AAA^ mutation *in vivo,* although rescuing of phenotypes with mito-RRM was not practical *in vivo.* The *in vivo* phenotypes therefore reflect the combined consequences of loss of RNA binding along with the sharp decrease in total Synj2 protein levels. Because Synj2 binds numerous mitochondrial transcripts including PINK1 and ATP51β(5), we examined mitochondrial respiration and ultrastructure. We employed synaptosomes, reasoning that these distal-most portions of neurons are most likely to depend on RNA transport for local protein synthesis. We isolated synaptosomes from 6-month-old WT and *Synj2*^AAA^ mice and performed Seahorse extracellular flux analysis to monitor oxygen consumption rates (Figure 3A).

**Figure 3.**
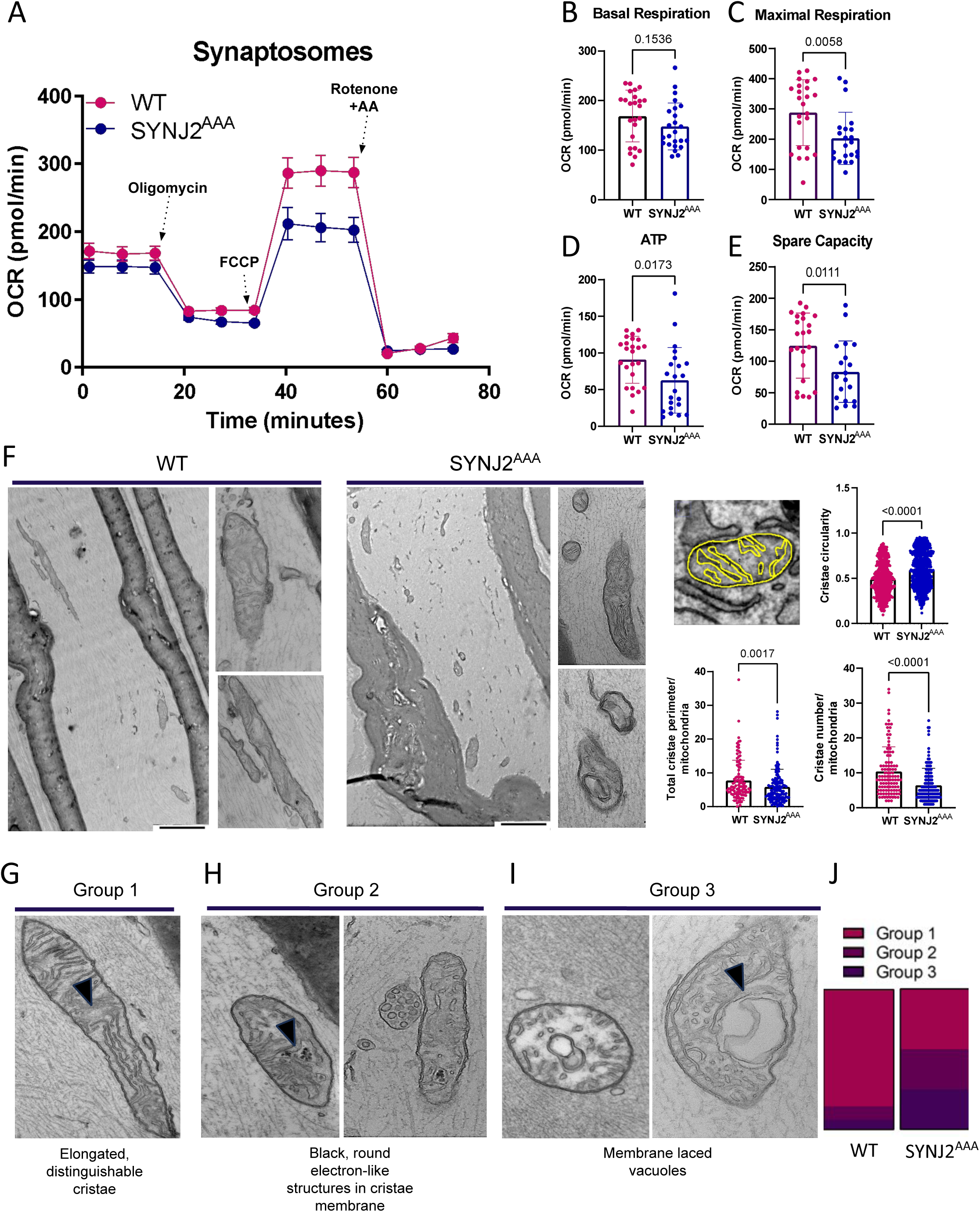
SYNJ2 RRM mutations impair mitochondrial respiration and ultrastructure. **(A)** Synaptosomes isolated from 6-month-old WT and Synj^AAA^ mouse brains were used to measure real-time oxygen consumption rate (OCR), measured at baseline followed by sequential injection of oligomycin (ATP synthase inhibitor), FCCP (mitochondrial uncoupler), and rotenone plus antimycin A (electron transport chain inhibitors) **B-E)** Quantification of basal respiration **(B)**, maximal respiration **(C)**, ATP production, and **(D)** spare respiratory capacity **(E)**. **n** = 20-24; error bars represent SD. **F)** Representative transmission electron micrographs of axonal mitochondria from WT and Synj2^AAA^ sciatic nerves. Quantitative analysis of cristae circularity, total mitochondrial number, mitochondrial perimeter, and cristae number per mitochondrion. **n** = 150-175 mitochondria. **G)** Representative electron micrographs of Class 1 **(G)**, Class 2 **(H)**, Class 3 **(I)** mitochondria. Class 1 mitochondria are elongated with well-organized, elongated cristae; Class 2 mitochondria have disorganized cristae with round, electron-dense deposits within the cristae; Class 3 mitochondria have disorganized cristae with membrane-laced vacuoles. Arrowheads indicate representative examples of each morphological category. (**J)** Quantification of the proportion of Class 1, Class 2 and Class 3 mitochondria in WT and SYNJ2^AAA^ sciatic nerve axons.

Synaptosomes were sequentially exposed to oligomycin, FCCP, and rotenone/antimycin A to assess parameters of mitochondrial respiration. While basal respiration was not convincingly different between genotypes (Figure 3B), maximal respiration (Figure 3C) was reduced in *Synj2*^AAA^ compared to WT controls. Additionally, both ATP production (Figure 3D) and spare respiratory capacity (Figure 3E) were decreased in *Synj2*^AAA^ synaptosomes, indicating compromised mitochondrial function in synaptic terminals.

To quantify mitochondrial ultrastructure, we examined longitudinal sections of axons in the sciatic nerve, which represent some of the longest axons in the body and therefore are where mRNA transport and local translation should be particularly important. Mitochondria were identified by electron microscopy based on the presence of a double membrane and internal cristae folds, distinguishing them from other axonal organelles. Electron microscopy revealed abnormalities in cristae morphology in *Synj2*^AAA^ mice. As a population, the Synj2^AAA^ mitochondria had less organized and less elongated cristae than in WT axons. *Synj2*^AAA^ mitochondria had fewer cristae per mitochondrion and the total cristae area per mitochondrion was decreased (Figure 3F).

Quantification revealed significant heterogeneity among *Synj2^A^*^AA^ mitochondria (Figure 3). Some looked normal, elongated and with well-formed cristae (Figure 3G). Others had abnormalities including round, electron-dense structures within the cristae (Figure 3H), greatly disordered cristae, or membrane-laced vacuoles (Figure 3I), which were identified as large, empty-appearing spaces bounded within the mitochondria. To better quantify the state of the mitochondria, we classified each mitochondrion into one of three categories: Class 1 were most normal with elongated mitochondria often aligned in parallel; Class 2 has electron dense deposits within cristae; Class 3 had less organized cristae and vacuoles among the disorganized cristae. Class 2 and 3 comprised 50% of the mitochondria in *Synj2*^AAA^ axons, but less than 20% in WT (Figure 3J). These ultrastructural defects are in accord with the observed decrease in maximal respiration in *Synj2*^AAA^ synaptosomes and potentially also with defective clearance of dysfunctional mitochondria by PINK1-dependent mitophagy. The presence of some mitochondria with normal morphology may explain why the functional deficits, though significant, are not more severe.

### Metabolic reprogramming in Synj2^AAA^ synaptosomes

To investigate the metabolic consequences of the *Synj2*^AAA^ mutation in synaptic terminals, we isolated synaptosomes from 6-month-old WT and *Synj2*^AAA^ mouse brains using discontinuous Ficoll density gradient centrifugation, extracted metabolites, and performed liquid chromatography-mass spectrometry (LC-MS/MS) to identify and quantify synaptic metabolites (Figure 4A). Principal component analysis of the resulting metabolomic profiles revealed clear separation between genotypes, with PC1 capturing 42.7% of total variance and PC2 capturing an additional 34.1%, together accounting for 77% of total metabolic variation and confirming robust, reproducible metabolic differences between WT and *Synj2*^AAA^ synaptosomes (Figure 4B). Volcano plot and hierarchical clustering analyses confirmed that metabolic changes were bidirectional, with altered metabolites distributed across both upregulated and downregulated categories (Figure 4C, D). Twenty metabolites of the detected 300 were significantly altered after correction for multiple comparisons (FDR < 0.05), with 15 elevated and 5 reduced in *Synj2*^AAA^ synaptosomes relative to WT controls. The most significant change was a reduction in cholesterol (p = 1.29 × 10⁻¹⁰, FDR = 3.73 × 10⁻⁸), a principal structural component of synaptic membranes required for lipid raft organization and synaptic vesicle release. This finding is consistent with the Synj2-associated RNA-IP dataset(5), in which transcripts encoding key cholesterol biosynthetic enzymes (DHCR7, NSDHL, ACAT2) were among those whose association with Synj2 was lost in *Synj2*^AAA^ neurons. Failed local translation of these biosynthetic enzymes may thus contribute directly to cholesterol depletion at synaptic terminals. Concurrently, acetylcarnitine was elevated (p = 1.24 × 10⁻⁶, FDR = 5.10 × 10⁻⁵), indicating that synaptic terminals are increasingly burning fatty acids for energy(10) rather than using lipids to maintain and build membranes.

**Figure 4.**
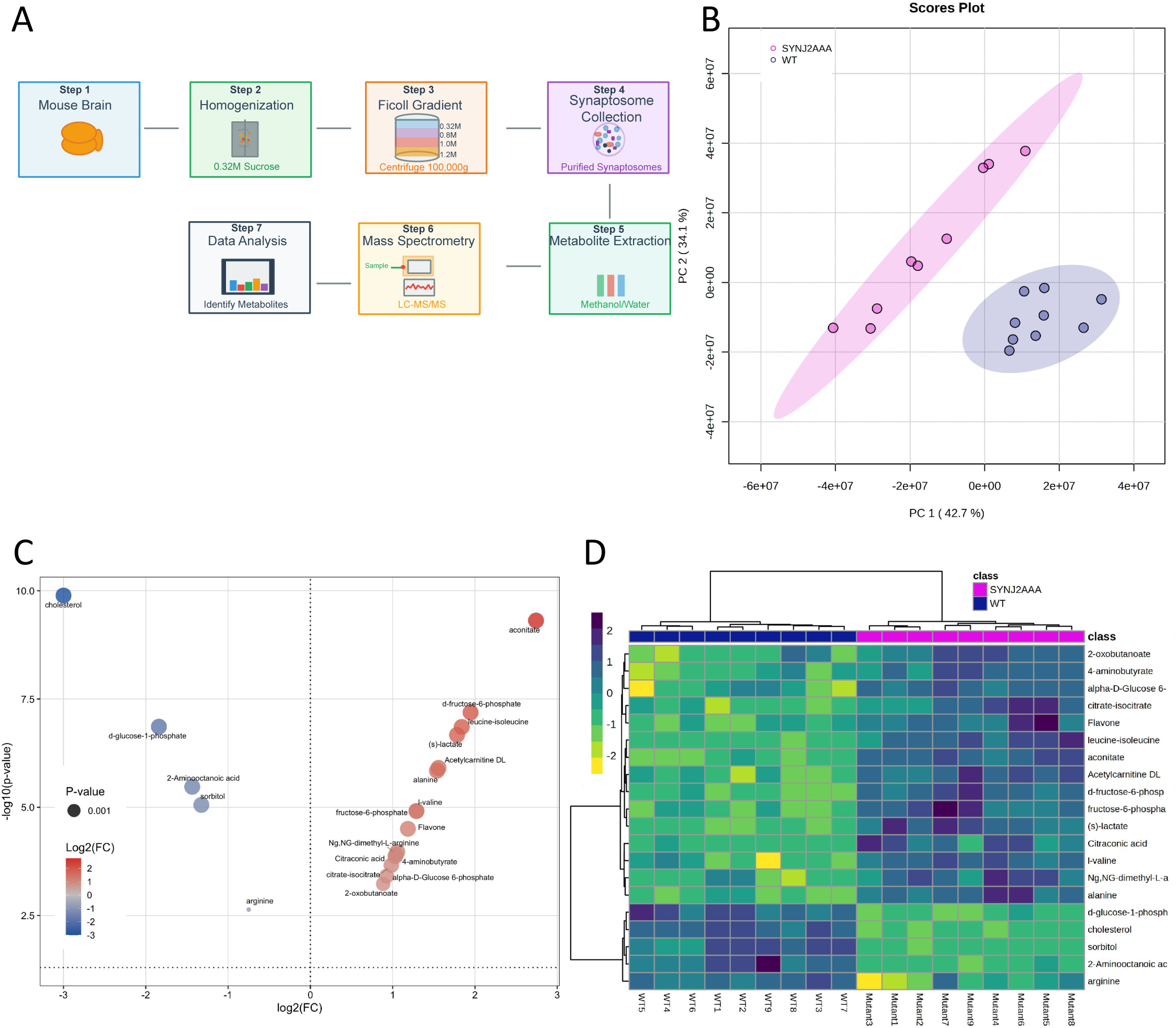
Metabolomic reprogramming in Synj2^AAA^ synaptosomes. **(A)** Experimental pipeline for synaptosome metabolomics. Synaptosomes were isolated from mouse brain by discontinuous Ficoll density gradient centrifugation, and metabolites were extracted using methanol/water and analyzed by liquid chromatography-tandem mass spectrometry (LC-MS/MS). **(B)** Principal component analysis of metabolomic profiles from WT and *Synj2*^AAA^ synaptosomes. PC1 captured 42.7% of total variance and PC2 captured 34.1%, together accounting for 77% of total metabolic variation. WT and *Synj2*^AAA^ samples form distinct, well-separated clusters, indicating robust and reproducible metabolic differences between genotypes. n = 9 biological replicates per genotype. **(C)** Volcano plot of significantly changed metabolites comparing *Synj2*^AAA^ to WT synaptosomes. Each circle represents one metabolite. Circle color indicates direction and magnitude of change, with blue representing decreased and orange/red representing increased metabolites in *Synj2*^AAA^ relative to WT; color intensity reflects log₂(fold change) on a continuous scale. Circle size is inversely proportional to p-value, with larger circles indicating greater statistical significance. The horizontal dotted line indicates the significance threshold (p = 0.05). The vertical dotted line indicates zero-fold change. **(D)** Hierarchical clustering heatmap of the 20 metabolites that were significantly altered between WT and *Synj2*^AAA^ synaptosomes (FDR < 0.05, unpaired two-tailed t-test with Benjamini-Hochberg correction). Color scale represents normalized metabolite abundance, ranging from dark purple (high) to yellow (low). Each column represents one biological replicate; each row represents one metabolite. WT and *Synj2*^AAA^ samples cluster into separate groups, consistent with the PCA separation shown in panel B. n = 9 biological replicates per genotype; error bars represent SD.

Within the TCA cycle, aconitate, the intermediate between citrate and isocitrate was the second most significantly altered metabolite overall (p = 4.89 × 10⁻¹⁰, FDR = 7.07 × 10⁻⁸), and citrate-isocitrate was also elevated (p = 2.19 × 10⁻⁴, FDR = 3.72 × 10⁻³). The co-accumulation of these adjacent TCA cycle intermediates indicates a bottleneck at the isocitrate dehydrogenase step, consistent with the impaired maximal respiration observed in *Synj2*^AAA^ synaptosomes (Figure 3A). Three branched-chain amino acids were elevated, leucine-isoleucine (p = 1.40 × 10⁻⁷, FDR = 8.07 × 10⁻⁶), valine (p = 1.22 × 10⁻⁵, FDR = 2.95 × 10⁻⁴), and the leucine catabolism intermediate citraconic acid (p = 1.37 × 10⁻⁴, FDR = 2.47 × 10⁻³), consistent with increased protein breakdown or reduced capacity to metabolize branched-chain amino acids under conditions of mitochondrial stress(11, 12). Additional markers of amino acid catabolism were increased, including alanine (p = 1.43 × 10⁻⁶, FDR = 5.17 × 10⁻⁵), 4-aminobutyrate (GABA; p = 1.06 × 10⁻⁴, FDR = 2.19 × 10⁻³), and 2-oxobutanoate (p = 5.88 × 10⁻⁴, FDR = 8.95 × 10⁻³). Arginine was reduced (p = 2.30 × 10⁻³, FDR = 0.033) while Ng,NG-dimethyl-L-arginine (ADMA), a molecule that blocks nitric oxide production by competing with arginine for the same enzyme, was concurrently elevated (p = 1.23 × 10⁻⁴, FDR = 2.37 × 10⁻³), suggesting that nitric oxide signaling may be impaired in *Synj2*^AAA^ synaptic terminals(13, 14). Seven additional metabolites showed modest reductions of uncertain individual significance after correction for multiple comparisons, including the nucleotide metabolites dAMP (p = 9.8 × 10⁻³) and dCMP (p = 0.020), citrulline (p = 0.010), S-adenosyl-L-homocysteine (p = 0.020), and glutamate (p = 0.038); while these should be interpreted with caution, their consistent direction of change aligns with the broader pattern of reduced biosynthetic capacity in *Synj2*^AAA^ synaptosomes.

### Upregulation of branched-chain amino acid and energy metabolism pathways in SYNJ2^AAA^ synaptosomes

To identify which biological pathways were most affected by the *Synj2*^AAA^ mutation, significantly altered metabolites were mapped onto known metabolic pathways using enrichment analysis (Figure 5A). The top enriched pathways included glycolysis/gluconeogenesis, the citrate cycle (TCA cycle), starch and sucrose metabolism, fructose and mannose metabolism, and alanine/aspartate/glutamate metabolism, all of which showed high enrichment ratios and strong statistical significance (Figure 5A). These results indicate that the metabolic changes in *Synj2*^AAA^ synaptosomes are not scattered randomly across metabolism but instead concentrate in interconnected energy-producing pathways.

**Figure 5.**
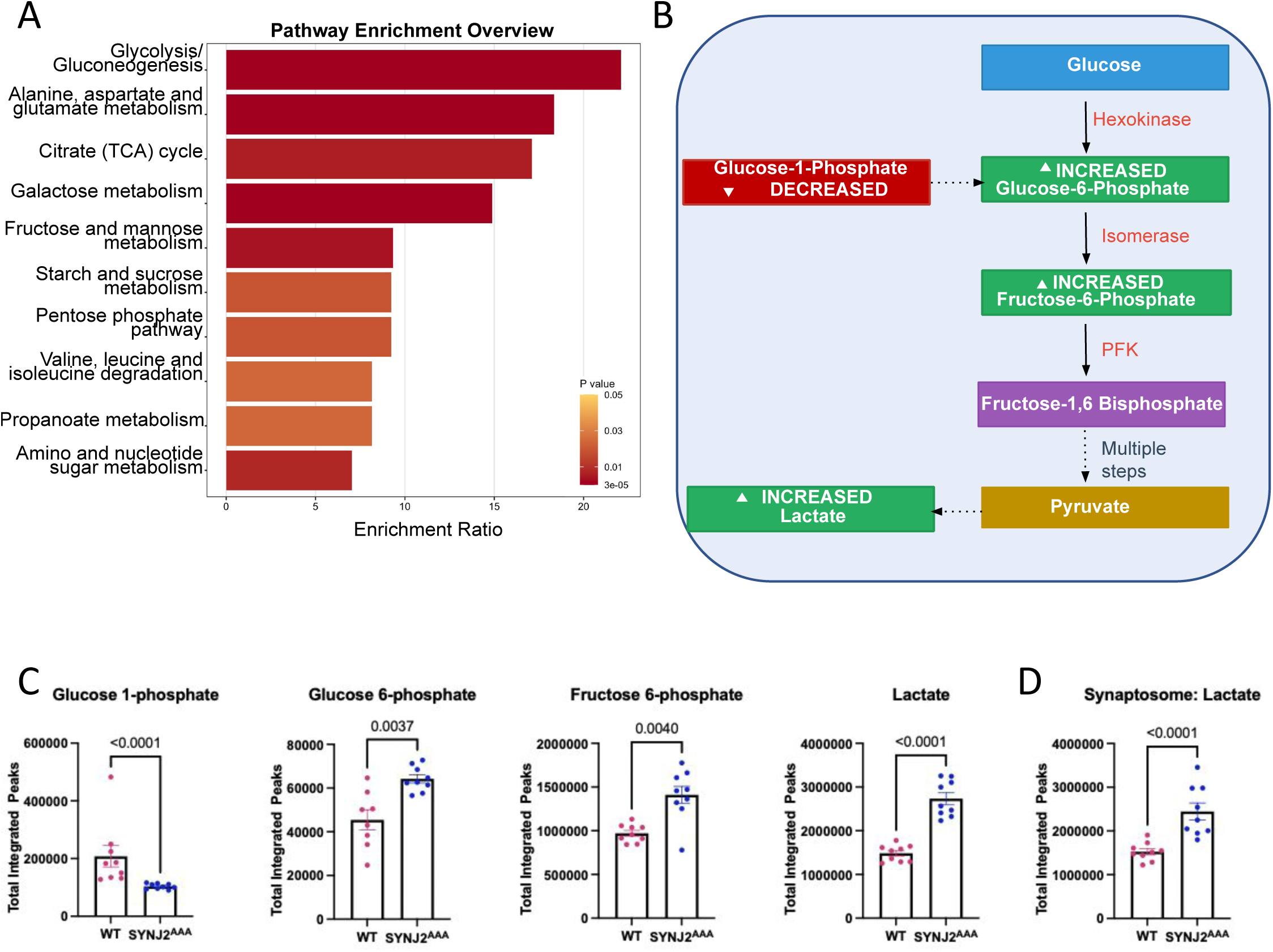
Upregulation of branched-chain amino acid and energy metabolism pathways in Synj2^AAA^ synaptosomes. **(A)** KEGG pathway enrichment analysis of significantly altered metabolites in *Synj2*^AAA^ synaptosomes, restricted to *Mus musculus* metabolic pathways. Bars represent enrichment ratio, calculated as the observed fraction of pathway metabolites altered relative to the fraction expected by chance. Bar color indicates statistical significance, with darker red reflecting lower p-values. **(B)** Schematic of the glycolysis pathway highlighting the direction of change for significantly altered metabolites in *Synj2*^AAA^ synaptosomes. Metabolites shown in green were elevated and metabolites shown in red were reduced relative to WT controls. **(C)** Quantification of individual glycolytic metabolite levels in WT and *Synj2*^AAA^ synaptosomes. Glucose-6-phosphate, fructose-6-phosphate, and lactate are elevated while glucose-1-phosphate is reduced. Statistical significance determined by unpaired two-tailed t-test. Error bars represent SD. n = 9 biological replicates per genotype. **(E)** Independent validation of lactate elevation in *Synj2*^AAA^ synaptosomes using a luminescence-based enzymatic assay (Lactate-Glo). Lactate levels are significantly higher in *Synj2*^AAA^ synaptosomes compared to WT controls, confirming the mass spectrometry finding through an orthogonal method. Statistical analysis by unpaired Student’s t-test. n =9 biological replicates per genotype; error bars represent SD.

To visualize how the significantly altered metabolites map onto the glycolytic pathway, we constructed a pathway diagram showing the direction of each change (Figure 5B), alongside quantification of the individual metabolites (Figure 5C). At the early steps of glycolysis, Synj2^AAA^ synaptosomes had higher glucose-6-phosphate (p = 0.0037) and fructose-6-phosphate (p = 0.0040) than wild-type synaptosomes. In contrast, glucose-1-phosphate was reduced (p < 0.0001). Glucose-1-phosphate is a branch-point metabolite that links glycolysis to glycogen metabolism. Lactate, the end product of anaerobic glycolysis, was elevated (p < 0.0001). Fructose-1,6-bisphosphate and pyruvate were not significantly altered. Taken together, higher early glycolytic intermediates and higher lactate are consistent with increased glycolytic flux in Synj2^AAA^ synaptic terminals. The reduction in glucose-1-phosphate is also consistent with depleted glycogen. Whether this reflects neuronal or glial glycogen pools within the synaptosome preparation remains unclear.

To independently validate the mass spectrometry finding that lactate was elevated in *Synj2*^AAA^ synaptosomes, we measured lactate levels in a separate synaptosome preparation using a luminescence-based enzymatic assay. Lactate levels were significantly higher in *Synj2*^AAA^ synaptosomes compared to WT controls (Figure 5D), confirming the mass spectrometry result through an orthogonal method and providing independent evidence that glycolytic flux is increased in *Synj2*^AAA^ synaptic terminals.

### Loss of RNA-binding by Synj2^AAA^ increases neuronal vulnerability to metabolic stress and excitotoxicity

To determine the consequences of Synj2 mutation on cellular viability, we asked how well neurons survived stresses. Primary mouse neurons were exposed to the mitochondrial uncoupler CCCP (10µM) for 6h and stained with 1 μM SYTOX Green to label dying cells and Hoechst dye to label nuclei. Cell death was significantly higher in *Synj2*^AAA^ neurons than in WT. (Figure 6A,B). Elevated levels of cleaved PARP in *Synj2*^AAA^ neurons after CCCP treatment indicated that the cell death occurred via caspase-mediated apoptosis (Figure 6C, D). Transduction with SYNJ2 mito-RRM, however, rescued both cell viability and cleaved PARP levels back to WT levels (Figure 6A-D), indicating that the RNA-binding function of SYNJ2 is critical for cellular resilience under this metabolic stress. To examine the vulnerability of neurons to excitotoxicity, we exposed EGFP-expressing primary cortical neurons to 100µM NMDA and 10µM glycine for 10 minutes and counted the percent of surviving GFP-positive neurons 24 h later. Synj2^AAA^ neurons were more sensitive to this excitotoxic stress (Figure 6E, F). As an additional measure of neuronal health, we measured neurite length in *Synj2*^AAA^ and WT iPSC-derived neurons that were placed in a glucose-free galactose medium, which forces reliance on oxidative metabolism by eliminating glucose-dependent glycolysis. Over five days in the galactose medium, *Synj2*^AAA^ neurons significantly retracted their neurites, which did not occur in WT neurons. SYNJ2^RRM^ treatment partially rescued neurite length, indicating that RNA localization to mitochondria could improve the metabolic flexibility needed to sustain the neurites, and that other properties of SYNJ2 likely also were needed (Figure 6G). Thus, SYNJ2^AAA^-induced metabolic reprogramming, while potentially compensatory in the short term and in the highly favorable conditions of standard culture medium, ultimately compromises neurons and leaves them vulnerable to metabolic and excitotoxic stresses.

**Figure 6.**
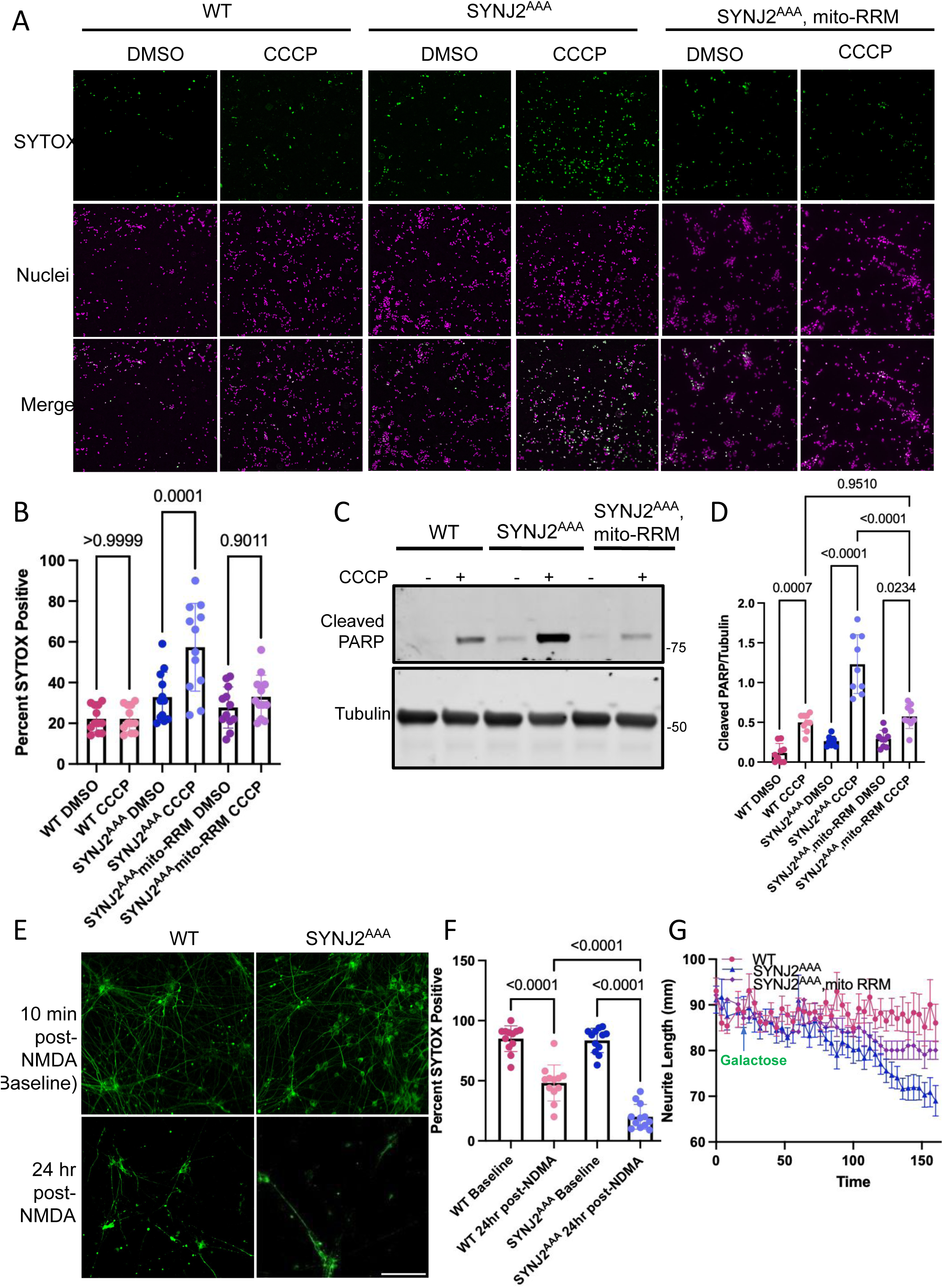
Loss of RNA-binding by Synj2^AAA^ increases neuronal vulnerability to metabolic stress and excitoxicity. **(A)** Cell viability of primary cortical mouse neurons (WT, SynjJ2^AAA^, and Synj2^AAA^ with mito-RRM expression) following a 6 hour exposure to 20 μM CCCP. Cell death was assessed using SYTOX staining (green) with nuclei (purple) counterstained. (**B)** Quantification shows percent SYTOX-positive (dead) cells relative to total cells. **n** = 12, error bars represent SD. (**C)** Immunoblot analysis of cleaved PARP (apoptosis marker) in WT, Synj2^AAA^, and Synj2^AAA^ neurons expressing mito-RRM. Neurons had a 6-hour exposure to 20 μM CCCP. β-tubulin serves as loading control. (**D)** Quantification shows relative cleaved PARP levels normalized to tubulin. **n** = 9, error bars represent SD. (**E)** Representative images of GFP-transfected WT and Synj2^AAA^ cortical neurons exposed to excitotoxic stress using NMDA treatment followed by washout after 10 minutes. **(F)** Quantification of GFP transfected WT and Synj2^AAA^ neurons following 10 min exposure of 100 uM NMDA and percentage of cells following 24 hr of NMDA exposure. **G)** Neurite outgrowth in WT, Synj2^AAA^ and Synj2^AAA^ neurons expressing mito-RRM under metabolic stress conditions. Culture medium was switched to 10 mM galactose with 0 mM glucose as indicated to force reliance on oxidative metabolism. Neurite outgrowth was measured over 6 days. **n** = 20-24; error bars represent SD.

## DISCUSSION

The results of this study build on our prior identification of Synj2a as an RNA-binding protein associated with mitochondria and the demonstration that it bound many transcripts encoding mitochondrial proteins. These included components of the electron transport chain, TCA cycle enzymes, and cholesterol biosynthesis pathway(5). Our metabolomic findings extend this framework from transcript-level observations to direct metabolic consequences in synaptic terminals and demonstrate that Synj2-dependent RNA transport has measurable downstream effects on synaptic energy metabolism and lipid composition. The breadth of disruption we observe, spanning mitochondrial quality control, respiratory capacity, ultrastructure, and synaptic metabolite profiles suggest that Synj2’s role extends far beyond synaptic vesicle recycling to encompass fundamental aspects of neuronal bioenergetics. Beyond the metabolic phenotypes, loss of Synj2 function renders neurons more vulnerable to mitochondrial and excitotoxic stress, connecting defects in local mRNA transport to neuronal survival under adverse conditions.

### RNA binding and the stability of the Synj2 protein

A three amino acid substitution in the RRM domain caused near-complete loss of full-length Synj2 protein from both mouse brain and cultured human iNeurons (Figure 1D). The drop in protein levels likely indicates that RNA-binding protects the protein from degradation. This parallels observations with other RNA-binding proteins, including HuR, TDP-43, and FMRP, whose stability depends on RNA association through mechanisms that include conformational protection, masking of degrons, and prevention of aggregation(17–19). For *Synj2^AAA^*, loss creates an interpretive challenge: although the amino acid changes did not themselves alter the enzymatic activity of the phosphatase domains (Figure 1B), the decreased protein levels could result in loss of both RNA transport and phosphatase activity. Our rescue experiments with the mitochondrially targeted RRM domain, which restores the RNA binding region but not the phosphatase domains, establish which phenotypes in cultured neurons are attributable specifically to RNA transport, while the remaining unrescued deficits may point to contributions from the phosphatase domains or other effects of Synj2 protein dosage. For phenotypes detected in vivo and in synaptosomes, though the importance of the phosphatase domains cannot be discounted, the correlation of defects with mRNA known to bind Synj2 strongly implicates the RNA-binding function as well.

Our demonstration that Synj2-mediated RNA localization directly impacts PINK1-dependent mitophagy in axons provides compelling evidence that mRNA transport to mitochondria has genuine functional consequences for neuronal quality control. We demonstrated that the RRM domain of Synj2 was required for: 1) the presence of the mRNA in distal regions, 2) the production of PINK1 protein in response to mitochondrial depolarization in axons, but not in the soma, and 3) the consequence of local PINK1 activation in the form of phosphoubiquitin on depolarized mitochondria. The restoration of phosphoubiquitin accumulation by restoring the RRM to mitochondria establishes a direct mechanistic link between RNA transport and mitochondrial homeostasis in distal neuronal compartments. The consequences of this disruption in a mitophagic pathway may contribute to other phenotypes we observed, including abnormal mitochondrial morphologies and greater sensitivity to neuronal stressors. The incomplete rescue of neuronal resilience to stresses, however, may reflect either inadequate RRM function when expressed outside the full-length Synj2 or the importance of Synj2 phosphatase domains. It is likely that all the domains of Synj2 contribute to mitochondrial function and thereby to neuronal resilience, albeit through distinct but complementary mechanisms. It may be surprising that the loss of so much Synj2 protein did not have a more drastic phenotype and impair the viability of the mice. The phosphatase activity of Synaptojanin 1 may account for this. With regard, however, to their RRM domains, Synj1 and Synj2 are not redundant; they differ in their ability to bind mRNAs, at least with regard to PINK1(5).

### Synj2 is important for synaptic metabolism and bioenergetics

The most statistically significant metabolic change in *Synj2*^AAA^ synaptosomes was a reduction in cholesterol, a finding that connects directly to the list of known mRNA species that bind Synj2(5), in which transcripts encoding key cholesterol biosynthesis enzymes were among those whose association with Synj2 was lost in Synj2^AAA^ protein. These include: DHCR7 (2.6-fold enrichment in WT over AAA), which catalyzes the final step of cholesterol biosynthesis; NSDHL (3.9-fold enrichment), a sterol dehydrogenase acting downstream of squalene; and ACAT2 (3-fold enrichment), which contributes to cholesterol esterification. Without proper localization of these transcripts to synaptic terminals, local synthesis of the enzymes that produce cholesterol is compromised, depleting a lipid that is essential for membrane integrity, lipid raft organization, and synaptic vesicle function(20–23). The concurrent elevation of acetylcarnitine further supports a shift away from lipid biosynthesis and toward lipid catabolism in *Synj2*^AAA^ terminals. It is important to note that while the RNA sequencing data from Harbauer et al. and our metabolomic findings together suggest this mechanism, direct validation including measurement of cholesterol biosynthesis enzyme activity, sterol flux experiments, and rescue of the cholesterol deficit by restoring individual biosynthetic transcripts will be needed to establish causality.

Within the TCA cycle, the co-accumulation of aconitate and citrate-isocitrate indicates a bottleneck at the isocitrate dehydrogenase step, consistent with the reduced maximal respiration in Seahorse analyses. Notably, IDH1 (3.4-fold enrichment) itself was among the Synj2-associated transcripts in the Harbauer et al. dataset of Synj2-associated mRNA, along with subunits spanning all five respiratory complexes including NDUFS1 and NDUFS2 (Complex I), SDHD and SDHA (Complex II), UQCRC1 (Complex III), COX5A (Complex IV), and ATP5F1B (Complex V), as well as the TCA cycle enzymes SUCLG1 and DLST. This breadth of representation across the respiratory chain suggests that loss of Synj2-dependent mRNA localization could compromise multiple steps of oxidative metabolism simultaneously, consistent with the global respiratory deficits observed here. The respiratory deficits and ultrastructural abnormalities reported here likely reflect failed local translation of multiple mitochondrial targets simultaneously.

### Metabolic compensation and its limits

The broader metabolic reprogramming in *Synj2*^AAA^ synaptosomes, elevated glycolytic intermediates, increased branched-chain amino acids, and upregulated amino acid catabolism, appears to represent a compensatory response to impaired oxidative metabolism. The elevation of branched-chain amino acids alongside their catabolism intermediate citraconic acid is best interpreted as increased protein breakdown or impaired degradation of branched-chain amino acids (BCAAs) rather than active biosynthesis(11, 12): mammals cannot synthesize these essential amino acids de novo, and the high enrichment ratio seen for the Kyoto Encyclopedia of Genes and Genomes (KEGG) biosynthesis pathway is a database artifact reflecting the co-annotation of all three BCAAs as terminal products of a non-mammalian pathway(15, 16). The mammalian-relevant signal comes from the degradation pathway, which showed both significant enrichment and a non-zero topological impact score, pointing to impaired catabolism as the biologically meaningful process. That said, the metabolomics data presented here are associative, and additional validation experiments including isotope tracing with ¹³C-labeled leucine or valine to directly measure BCAA flux, and enzymatic assays of BCAA catabolism pathway activity will be required to definitively establish whether impaired degradation or increased protein turnover drives the accumulation.

The reduction in glucose-1-phosphate alongside elevated downstream glycolytic intermediates could reflect depletion of glycogen stores in terminals that have been chronically upregulating glycolysis in vivo. However, neuronal glycogen is limited, and astrocytic glycogen is more abundant; since glial processes are a recognized contaminant of synaptosome preparations, whether this signal reflects neuronal or glial glycogen pools cannot be determined from these data alone. Cell-type-specific metabolic labeling experiments would be needed to resolve this question. Despite these compensatory metabolic shifts, functional assays reveal their limits.

Increased vulnerability to CCCP, NMDA excitotoxicity, and galactose-induced metabolic stress demonstrates that apparent metabolic activation masks underlying cellular fragility. These short-term adaptations that sustain basic function under standard conditions cannot substitute for the loss of mitochondrial quality control and local protein synthesis when neurons face additional stress.

### Implications for neurodegeneration

PINK1 mutations cause early-onset Parkinson’s disease(7), and our findings raise the possibility that impaired PINK1 mRNA localization could contribute to neuronal dysfunction even when the PINK1 coding sequence is intact, a form of localization pathology that extends beyond traditional loss-of-function mechanisms. Defects in mitochondrial transport have been described in many neurodegenerative diseases(24–27). If transport on mitochondria is a major means of supplying certain transcripts, such as PINK1, to distal axons, diminished local protein synthesis may exacerbate the consequences of poor mitochondrial transport. The cholesterol depletion we observe adds a further dimension to this vulnerability: age-related declines in brain cholesterol biosynthesis are well established(28–30) Neurons already operating with reduced synaptic cholesterol due to Synj2 dysfunction may be particularly susceptible to this decline, potentially contributing to the selective vulnerability of neuronal populations in neurodegenerative diseases where both mitochondrial dysfunction and lipid dysregulation are implicated. The metabolic inflexibility demonstrated in galactose medium is also particularly relevant to aging, where mitochondrial dysfunction progressively limits oxidative capacity and neurons must increasingly rely on compensatory metabolic strategies to survive.

More broadly, the synaptosomal metabolic reprogramming we observe mirrors signatures reported in degenerating neurons in Parkinson’s disease and other neurodegenerative conditions: suppressed oxidative phosphorylation, compensatory glycolytic upregulation, disrupted branched-chain amino acid catabolism, and impaired nitric oxide signaling at synaptic terminals (31–36). These convergences suggest that loss of Synj2-mediated mRNA transport to axonal mitochondria may recapitulate key features of the metabolic failure that precedes neuronal death in disease. The metabolic inflexibility we observe in galactose medium is particularly relevant to aging, when mitochondrial function progressively declines and neurons must rely increasingly on compensatory strategies to survive. Neurons already compromised by Synj2 dysfunction or defective mitochondrial transport of Synj2 and its associated mRNA may be disproportionately vulnerable to this age-associated energetic stress. This vulnerability is compounded by the lipid dysregulation we describe, since age-related declines in brain cholesterol biosynthesis are well established (23, 28), and neurons with reduced synaptic cholesterol due to disrupted Synj2-dependent mRNA transport may be especially susceptible to it.

Together, these findings establish Synj2 as a link between mRNA localization, mitophagy, lipid metabolism, and mitochondrial bioenergetics in neurons, providing direct metabolic evidence for the functional significance of the RNA transport activity first described by Harbauer et al(5). Synj2 deficiency simultaneously depletes synaptic cholesterol and forces compensatory reliance on glycolysis and amino acid catabolism, while leaving neurons more vulnerable to stress. Thus, the RNA transport and lipid phosphatases function jointly to maintain healthy synaptic terminals. mRNA localization to synaptic mitochondria is likely to be important for the transcriptional and translation regulation that makes neurons resilient. That resilience is particularly important in the context of aging and neurodegeneration, where mitochondrial function and lipid metabolism are both progressively compromised.

## METHODS AND MATERIALS

### Site-directed mutagenesis

Synj2 mutant constructs were generated by site-directed mutagenesis using the Q5 site-directed mutagenesis kit (NEB) according to the manufacturer’s instructions. The starting template was a plasmid encoding Flag-tagged mitochondrial-localized SYNJ2a. The following mutations were introduced: RNA binding deficient mutant (SYNJ2^AAA^) containing V892A, Q934A, and L936A substitutions; Sac1 domain inactive mutant (SYNJ2^D388N^) containing the D388N substitution; and phosphatase inactive mutant (SYNJ2^N538K^) containing the N538K substitution. Complementary primer pairs were designed to introduce each mutation with at least 15 bases flanking the mutation site. PCR reactions were performed using Q5 High Fidelity DNA polymerase. Following PCR, the parental methylated DNA was digested, phosphorylation and ligation were performed using the KLD enzyme Mix (NEB) for 5 min at 25°C, and the reaction products were transformed into XL10-Gold ultracompetent cells. All mutations were confirmed by DNA sequencing of the entire SYNJ2 coding region to ensure no unintended mutations were introduced.

### Generation of SYNJ2^AAA^ Knock-in Mice using CRISPR/Cas9

The BCH Gene Manipulation Core generated the SYNJ2^AAA^ mice that introduced V892A, Q934A, and L936A mutations into the SYNJ2 gene by using CRISPR/Cas9 genome editing. Guide RNAs (gRNAs) targeted exon 22 of the mouse *Synj2* gene. The gRNA sequences were: 5’-CTGGAGGGTTACAACAACGG-3’ and 5’-ACGTCACCAGCATCTGCCCT-3’ (targeting the Q934/L936 region). A single-stranded DNA oligonucleotide (ssODN) repair template was designed to introduce all three-point mutations simultaneously while maintaining silent mutations to prevent re-cutting by Cas9. The 1116-nucleotide ssODN contained the desired mutations flanked by nucleotides homology on each side. Cas9 protein, gRNAs, and the ssODN repair template were co-injected into fertilized C57BL/6N oocytes by pronuclear injection. Injected embryos were transferred into pseudopregnant ICR foster mothers, and resulting offspring were genotyped by PCR amplification of the target region followed by Sanger sequencing. Founder mice carrying the desired mutations were bred to homozygosity.

### Animal care and use

All animal procedures were approved by Boston Children’s Hospital IACUC and performed in accordance with the National Institutes of Health Guide for the Care and Use of Laboratory Animals. Mice were housed under a 12hr light/dark cycle with ad libitum access to food and water in housing conditions at Boston Children’s Hospital. Mice were allocated to experimental groups by genotype and littermates were used as matched controls wherever possible. Image based quantification were blinded to genotype during image acquisition and analysis.

### Mouse genotyping

Mouse genotyping was performed using tail biopsy samples collected at weaning. Tail snips were obtained using sterile scissors and placed in individual 1.5mL tubes. Genomic DNA was extracted using DirectPCR lysis reagent (Viagen Biotech) according to manufacturer instructions. The resulting DNA was used for PCR amplification using primers flanking the SYNJ2^AAA^ mutation sites in exon 22. Forward primer (5’-TTGGCCACTCTGTGCTAAGG-3’) and reverse primer (5’-GTGTGAACAGTGTCCCCCATAAC-3’) were used to amplify a 1190 bp fragment containing all three mutation sites (V892A, Q934A, L936A). PCR reactions were performed with Q5 high fidelity DNA polymerase. Products were analyzed by agarose gel electrophoresis. For definitive genotype determination, PCR products were purified using the QIAquick PCR Purification Kit (Qiagen) and subjected to Sanger sequencing.

### Generation of SYNJ2^AAA^ iPSCs using CRISPR/Cas9

Human induced pluripotent stem cells (iPSCs) were cultured on Matrigel-coated plates in StemFlex medium (STEMCELL Technologies) at 37°C with 5% CO₂. For CRISPR/Cas9 editing, iPSCs were dissociated using Accutase and resuspended at 1×10⁶ cells/mL in electroporation buffer. The same gRNA sequence and ssODN repair template described above were adapted for human SYNJ2 with appropriate codon optimization. The gRNA sequence was cloned into a Cas9-encoded plasmid (pSpCas9(BB)-2A-Puro (PX459) V2.0, Addgene) and was electroporated into 2×10⁶ iPSCs with the ssODN containing the mutations using the Lonza Amaxa Nucleofector (Lonza). Following electroporation, cells were plated on Matrigel in mTeSR1 medium supplemented with 10 μM Y-27632 ROCK inhibitor. After 48 hours, cells were plated one cell per well in a 96 well plate. Individual colonies were picked after 10-14 days and expanded for genotyping. Genomic DNA was extracted using QuickExtract DNA Extraction Solution, and the target region was amplified by PCR using primers flanking the editing site. Positive clones were confirmed by Sanger sequencing. Wild-type cells showed the original amino acid sequences (GTG for V892, CAG for Q934, CTG for L936) and homozygous SYNJ2^AAA^ cells showed mutant sequences (GCC for A892, GCC for A934, GCC for A936).

### Primary cortical neuron isolation and maintenance

Primary cortical neurons were isolated from postnatal day 0 (P0) WT and *SYNJ2*^AAA^ mutant mouse pups. Pups were sacrificed by decapitation and brains were rapidly dissected in ice-cold Hanks’ Balanced Salt Solution (HBSS) without calcium and magnesium. Cortices were carefully dissected under a stereomicroscope, removing meninges and subcortical structures, and transferred to fresh ice-cold HBSS. Cortical tissue was digested with 0.25% trypsin-EDTA at 37°C for 5 minutes with gentle agitation every 2 minutes. Trypsin activity was terminated by adding an equal volume of DMEM supplemented with 10% fetal bovine serum and 1% penicillin/streptomycin. Dissociated tissue was centrifuged at 200 × g for 5 minutes at room temperature and the pellet was resuspended in Neurobasal medium supplemented with 2% B27, 0.5 mM GlutaMAX, and 1% penicillin/streptomycin. Cells were counted using a hemocytometer and viability was assessed by trypan blue exclusion. Primary cortical neurons were plated on poly-D-lysine coated coverslips or culture dishes in a humidified incubator at 37°C with 5% CO₂. After 1hr to allow cell attachment, the medium was replaced with fresh Neurobasal medium supplemented with 2% B27, 0.5 mM GlutaMAX, and 1% penicillin/streptomycin. To inhibit glial proliferation, cytosine arabinoside (AraC) was added to the culture medium on days in vitro (DIV) 2-3. Half of the culture medium was replaced every 3-4 days with fresh Neurobasal/B27 medium without AraC. Neurons were maintained for 7-14 days in vitro to achieve synaptic maturation before experimental use.

### iPSC-derived cortical neuron differentiation

Human iPSCs (WT and SYNJ2^AAA^) were differentiated into cortical neurons via NGN2 overexpression using a tetracycline-inducible system. iPSCs were maintained on Matrigel-coated plates in StemFlex medium until 70-80% confluent then transduced with lentiviral vectors encoding pTet-O-NGN2-puromycin and rtTA in mTeSR1 medium. After 24 hours, viral medium was replaced with fresh StemFlex medium. NGN2 expression was induced by culturing cells for 24hrs in KSR media (KO DMEM supplemented with Knockout serum replacement, 1x non-essential amino acids, 10 μM β-mercaptoethanol, 2 mM Glutamax) containing doxycycline (2 μg/mL). Cells were then selected with puromycin (1ug/mL) in 1:1 mixture of KSR media medium and N2B medium (DMEM/F12 supplemented with 1× non-essential amino acids, 2mM GlutaMAX, N2 supplement). After 24 hours of doxycycline induction, medium was switched to N2B medium supplemented with 1× B27 supplement. Doxycycline (2 μg/mL) was maintained throughout the 4-day differentiation period with daily medium changes. By day 4, cells displayed mature neuronal morphology with extensive neurite outgrowth and were either harvested for analysis or maintained for downstream functional studies. For extended cultures, neurons were maintained for 14-21 days in Neurobasal medium supplemented GDNF (10ng/mL, Thermo), CNTF(10ng/mL, Thermo), BDNF (10ng/mL, Thermo), B27 (1:100, Thermo), doxycycline (2ug/mL, Sigma) and puromycin (10ug/mL, Gibco). Doxycycline was withdrawn at DIV10 to eliminate its inhibitory effect on mitochondrial protein synthesis.

### SYNJ2 Phosphatase Assay

Flag-tagged full-length SYNJ2 constructs: WT, RNA-binding-deficient (SYNJ2^AAA^), Sac1 domain-inactive (SYNJ2^D388N^), and phosphatase-inactive (SYNJ2^N538K^) were transiently expressed in HEK293 cells by transfection with polyethyleneimine (PEI) at a 3:1 PEI:DNA ratio. Briefly, DNA and PEI were combined in 1 mL serum-free DMEM, incubated for 15 min at room temperature, and added dropwise to cells. At the time of harvest, cell pellets were resuspended in lysis buffer (1% NP-40, 4mM MgCl_2_, 150mM NaCl, 25mM Tris, supplemented with protease inhibitor at 1:200 and 1mM PMSF) and lysed on ice for 20 min.

Lysates were clarified by centrifugation (13,000 rpm, 30 min, 4°C), and a 50 μL aliquot of the supernatant was reserved as input. Flag-tagged SYNJ2 was immunoprecipitated by rotating with anti-FLAG beads (Sigma) for 2 hr at 4°C. Beads were pelleted by centrifugation (5,000 xg, 2 min, 4°C) and a flowthrough aliquot was collected before the beads were washed twice in lysis buffer and twice in assay buffer (30 mM HEPES, 100 mM KCl, 1 mM EGTA, 2 mM MgCl2). Washed beads were resuspended in 60 μL assay buffer. Phosphatase activity was measured using a malachite green-based assay. Bead-bound SYNJ2 was incubated with 3 μg PIPs substrate (Fisher) for 30 min at 37°C in a total volume of 60 μL, after which beads were pelleted by centrifugation (5,000 rcf, 30 sec, 4°C) and the supernatant containing released free phosphate was collected. Malachite green phosphate assay kit (Sigma) and phosphate standards were prepared according to the manufacturer’s instructions. Supernatant was added to a 96-well plate followed by malachite green reagent, mixed gently, and absorbance was read immediately and again after a 30 min incubation at room temperature at 620 nm. n=12 for each group.

### Quantitative real-time PCR

RNA was isolated from synaptosomes prepared from six month old WT and SYNJ2^AAA^ mice (see synaptosome isolation from mouse brain) and from whole brain tissue WT and SYNJ2^AAA^ mice using TRIzol reagent, according to the manufacturer’s instructions. RNA concentration and purity (A260/280) was determined using a NanoDrop spectrophotometer. Complementary DNA (cDNA) was synthesized from 1ug total RNA using the iScript cDNA Synthesis Kit (Bio-Rad) according to the manufacturer’s instructions. Quantitative PCR was performed on a CFX96 Real-Time PCR System, Biorad using SYBR Supermix. For synaptosome samples, primers targeting Pink1, ND1, ATP5β and β-actin were used and tubulin served as the reference gene. For whole brain samples, primers targeting total Synj2 and the mitochondrial-specific Synj2a isoform (amplifying the mitochondrial-targeting sequence) were used to assess relative transcript abundance between genotypes, and were normalized to Tubulin. Relative gene expression was calculated using the 2^(-ΔΔCt) method. Data were collected from three independent synaptosome preparations or brain samples per genotype, and statistical comparisons between WT and SYNJ2^AAA^ were performed using unpaired two-tailed Student’s t-tests. n=9 for each group.

### Protein isolation and immunoblot analysis

WT SYNJ2 and SYNJ2^AAA^ mutant mice at 6 months were sacrificed by CO₂ asphyxiation followed by cervical dislocation, and brains were rapidly dissected and homogenized in ice-cold lysis buffer. iPSC-derived neurons were differentiated for 18-21 days. Cells were washed twice with ice-cold PBS and lysed directly in the same lysis buffer used for brain tissue (150 mM NaCl, 5 mM EDTA, 1% NP-40, 0.5% sodium deoxycholate, 0.1% SDS, 50 mM Tris) supplemented with protease inhibitor cocktail (Roche). Brain tissue was homogenized using a Dounce homogenizer (20 strokes) on ice and sonicated for 10 sec; iPSC-derived neuron lysates were not sonicated. All lysates were incubated on ice for 30 min and clarified by centrifugation at 14,000 × g for 15 min at 4°C. Supernatants were collected, and protein concentration was determined by BCA assay. For immunoblotting of SYNJ2, samples were instead prepared in a separate lysis buffer (150 mM NaCl, 1% NP-40, 25 mM Tris, 4 mM MgCl^2^), mixed with 6X Laemmli buffer plus 5% β-mercaptoethanol, and heated at 50°C for 5 min prior to loading. All other samples were mixed with 6X Laemmli buffer plus 5% β-mercaptoethanol and boiled at 95°C for 5 min.

For immunoblot analysis, equal amounts of protein (20–30 μg, 50ug for SYNJ2) were separated by SDS-PAGE on 8% acrylamide gels and transferred to PVDF membranes. Membranes were blocked in 5% non-fat dry milk in PBS-T (0.1% Tween-20) for 1 hr at room temperature, then incubated with primary antibodies overnight at 4°C. After three washes in PBS-T, membranes were incubated with HRP-conjugated secondary antibodies (1:10,000) for 2 hr at room temperature. Proteins were visualized by LI-COR and quantified by densitometry using ImageJ software. To assess apoptosis following mitochondrial dysfunction, primary cortical and iPSC-derived neurons were treated with 10uM CCCP or DMSO vehicle (0.1%) for 6hrs, then washed and lysed as above. Cleaved PARP was detected using rabbit anti-cleaved PARP) and mouse anti-tubulin as loading control, applied overnight at 4C, followed by secondary antibodies. n=9 for each group

### Immunofluorescence

For immunofluorescence analysis, both primary cortical neurons and iPSC-derived neurons were plated on 18 mm glass coverslips pre-coated with poly-D-lysine (50 μg/mL) for primary neurons or poly-L-ornithine (10 μg/mL) plus laminin (10 μg/mL) for iPSC-derived neurons. Neurons were fixed with 4% paraformaldehyde in PBS for 5 minutes at 37C, followed by three washes with PBS. Fixed cells were permeabilized with 0.1% Triton X-100 in PBS for 10 minutes and blocked with 5% normal goat serum plus 1% bovine serum albumin in PBS for 1 hour at room temperature. Primary antibodies were applied overnight at 4°C. After washing three times with PBS, coverslips were incubated with appropriate Alexa Fluor-conjugated secondary antibodies (1:1000, Thermo Fisher Scientific) for 2 hours at room temperature in the dark. Nuclei were counterstained with DAPI (1 μg/mL) for 5 minutes, and coverslips were mounted on glass slides using ProLong Gold antifade mounting medium. Images were acquired using a fluorescence microscope with a 20× objective lens. For each coverslip, at least 10 random fields were captured, and total cell number (DAPI-positive nuclei) and neuronal marker-positive cells (NeuN and MAP2) were quantified using ImageJ software. This analysis confirmed that neuronal differentiation purity and efficiency were comparable between WT and SYNJ2^AAA^ conditions for both primary cortical neurons (>95% NeuN-positive, >90% MAP2-positive) and iPSC-derived neurons (>85% NeuN-positive, >80% MAP2-positive), validating that genetic modification did not affect neuronal differentiation capacity. n=9-12 for each group.

### RNAscope in situ hybridization and immunofluorescence

For combined RNAscope and immunofluorescence analysis, both primary cortical neurons and iPSC-derived neurons were plated on 35 mm glass bottom dishes with 10 mm micro-well #1 cover glass (Fisher) pre-coated with poly-L-lysine for 2 hours at room temperature, followed by three washes with sterile water and air drying. RNAscope in situ hybridization was performed using the RNAscope® 2.5 HD Detection Reagent kit (Advanced Cell Diagnostics) according to manufacturer’s instructions with minor modifications for cultured cells. Briefly, neurons were fixed with 4% paraformaldehyde in PBS for 5 minutes at 37C, followed by dehydration serially in 50%, 70%, and 100% ethanol for 1 min each at room temperature. Cells were treated with Protease Plus for 10 minutes at room temperature, then washed twice with distilled water. The PINK1 probe (catalog #314541, Advanced Cell Diagnostics) was hybridized to target mRNA by incubating at 40°C for 2 hours in a HybEZ Hybridization System. Following probe hybridization, amplification was performed and PINK1 mRNA signals were detected using Red-B/Red-A according to kit instructions. Following RNAscope detection, immunofluorescence was performed to visualize mitochondria. Cells were blocked with 5% normal goat serum plus 1% bovine serum albumin in PBS for 1 hour at room temperature. Primary antibody against ATP5A (1:500, Abcam #ab14748) was applied overnight at 4°C to label mitochondria. After three washes with PBS, cells were incubated with Alexa Fluor 488-conjugated goat anti-mouse secondary antibody (1:1000, Thermo Fisher Scientific) for 2 hours at room temperature in the dark. Nuclei were counterstained with DAPI (1 μg/mL) for 5 minutes. Cells were briefly washed with PBS and maintained in PBS for immediate imaging. Images were acquired using a confocal microscope (Zeiss LSM 880) with a 63× oil immersion objective lens. For each dish, at least 10-15 neurons were imaged using sequential acquisition to avoid bleed-through between channels. PINK1 mRNA puncta were quantified within ATP5A-positive mitochondrial regions using ImageJ with the colocalization threshold plugin. Data were collected from at least three independent experiments for each cell type and genotype with colocalization analysis performed to determine the spatial relationship between PINK1 mRNA and ATP5A.n=12-15 for each group.

### Synaptosome isolation from mouse brain

Synaptosomes were isolated from fresh mouse brain tissue by discontinuous Ficoll density gradient centrifugation. Six-month-old WT and SYNJ2^AAA^ mice were sacrificed by CO₂ asphyxiation followed by cervical dislocation. Ficoll gradients were prepared, one per animal, layering 4 mL 13% Ficoll, 1 mL 9% Ficoll, and 4 mL 6% Ficoll from bottom to top. Brains were excised, and white matter was removed to the extent possible before homogenization in 30 mL ice-cold homogenization buffer supplemented with 30 μL PMSF and 30 μL pepstatin A using a glass-Teflon homogenizer (20 strokes). The homogenate was centrifuged at 5,000 rpm for 2 min (SS34 rotor, Sorvall) to pellet nuclei and cell debris, and the resulting supernatant (S1) was collected and centrifuged at 11,000 rpm for 12 min (SS34 rotor) to yield pellet P2. P2 was resuspended in 18 mL homogenization buffer, avoiding the dark brown mitochondria-enriched fraction, and 3 mL aliquots of the resuspended pellet were loaded onto each Ficoll gradient using a pipette with the tip end cut off to minimize shearing damage to synaptosomes. Gradients were centrifuged at 22,500 rpm for 35 min (SW41 rotor, Beckman), and synaptosomes were collected from the 9–13% Ficoll interface using a Pasteur pipette and transferred to a fresh SS34 tube. Synaptosomes were washed by adding 30 mL homogenization buffer and centrifuging at 11,000 rpm for 12 min (SS34 rotor); the final pellet (P3) was resuspended in 2.5 mL homogenization buffer per brain, supplemented with PMSF and pepstatin A.

### PINK1 protein immunofluorescence

For PINK1 protein analysis, primary cortical neurons and iPSC-derived neurons were plated on poly-L-lysine coated coverslips and maintained as described above. To induce mitochondrial damage and PINK1 accumulation, neurons were treated with carbonyl cyanide m-chlorophenyl hydrazone (CCCP, 10 μM, Sigma-Aldrich) in culture medium for 3 hours at 37°C. Control neurons received an equivalent volume of DMSO vehicle (0.1% final concentration). Following treatment, neurons were immediately fixed with 4% paraformaldehyde in PBS for 5 minutes at 37C and washed three times with PBS. Fixed cells were permeabilized with 0.1% Triton X-100 in PBS for 5 minutes and blocked with 5% normal goat serum plus 1% bovine serum albumin in PBS for 1 hour at room temperature. Primary antibodies were applied overnight at 4°C: rabbit anti-PINK1 (1:200, Novus Biologicals) and mouse anti-ATP5A (1:500, Abcam) for mitochondrial labeling. After washing three times with PBS, coverslips were incubated with appropriate Alexa Fluor-conjugated secondary antibodies: Alexa Fluor 568 goat anti-rabbit (1:1000) for PINK1 detection and Alexa Fluor 488 goat anti-mouse (1:1000) for ATP5A, for 1 hour at room temperature in the dark. Nuclei were counterstained with DAPI (1 μg/mL) for 5 minutes, and coverslips were mounted using ProLong Gold antifade mounting medium. Images were acquired using a confocal microscope (Zeiss LSM 880) with a 63× oil immersion objective lens. For each condition, at least 15-20 neurons were imaged using sequential acquisition. PINK1 fluorescence intensity was quantified within ATP5A-positive mitochondrial regions using ImageJ. Colocalization analysis between PINK1 and ATP5A was performed using the colocalization threshold plugin, and data were expressed as colocalization coefficients (Manders’ coefficients). Background fluorescence was subtracted using regions of interest from cell-free areas. Data were collected from at least three independent experiments for each condition, with quantitative analysis comparing CCCP-treated versus vehicle-treated controls to assess PINK1 recruitment to damaged mitochondria. n=12-15 for each group.

### Mitochondrial ubiquitination immunofluorescence

For detection of mitochondrial ubiquitination, primary cortical neurons and iPSC-derived neurons were plated on poly-L-lysine coated coverslips and maintained as described above. To induce mitochondrial dysfunction and subsequent ubiquitination, neurons were treated with antimycin A (10 μM, Sigma-Aldrich) dissolved in DMSO for 6 hours at 37°C in culture medium. Control neurons received an equivalent volume of DMSO vehicle (0.1% final concentration). The 6-hour treatment period was chosen to allow sufficient time for mitochondrial damage recognition and ubiquitin conjugation while maintaining cell viability. Following antimycin A treatment, neurons were immediately fixed with 4% paraformaldehyde in PBS for 5 minutes at room temperature and washed three times with PBS. Fixed cells were permeabilized with 0.1% Triton X-100 in PBS for 5 minutes and blocked with 5% normal goat serum plus 1% bovine serum albumin in PBS for 1 hour at room temperature to prevent non-specific antibody binding.

Primary antibodies were applied overnight at 4°C: mouse anti-phospho-ubiquitin (Ser65, 1:500, Cell Signaling Technology #62802) which specifically recognizes PINK1-phosphorylated ubiquitin, and rabbit anti-ATP5A (1:500, Abcam #ab14748) for mitochondrial identification. After washing three times with PBS, coverslips were incubated with secondary antibodies: Alexa Fluor 568 goat anti-mouse (1:1000) for phospho-ubiquitin detection and Alexa Fluor 488 goat anti-rabbit (1:1000) for ATP5A, for 1 hour at room temperature in the dark. Nuclei were counterstained with DAPI (1 μg/mL) for 5 minutes, and coverslips were mounted using ProLong Gold antifade mounting medium. Images were acquired using a confocal microscope (Zeiss LSM 880) with a 63× oil immersion objective lens. For each condition, at least 15-20 neurons were imaged using sequential acquisition to prevent channel bleed-through. Mitochondrial phospho-ubiquitin marking was quantified by measuring colocalization between phospho-ubiquitin (Ser65) and ATP5A signals using ImageJ with the colocalization threshold plugin. Colocalization analysis was performed using Manders’ coefficients, with results expressed as the fraction of ATP5A-positive mitochondria that were also phospho-ubiquitin-positive. Background fluorescence was subtracted using regions of interest from cell-free areas. Data were collected from at least three independent experiments for each genotype, with quantitative analysis comparing antimycin A-treated versus vehicle-treated controls. n=12-15 for each group.

### Seahorse mitochondrial stress test

Mitochondrial respiratory function in synaptosomes was assessed using the Seahorse XFe96 Extracellular Flux Analyzer (Agilent Technologies) with the Seahorse XF Cell Mito Stress Test Kit. Fresh synaptosomes isolated from WT and SYNJ2^AAA^ mice were used immediately following the Ficoll gradient purification described above. Synaptosome protein concentration was adjusted to 25 μg per well in Seahorse XF Base Medium supplemented with 10 mM glucose, 1 mM pyruvate, and 2 mM L-glutamine, with pH adjusted to 7.4 at 37°C. Seahorse XFe96 cell culture microplates were pre-coated with Cell-Tak cell adhesive for 20 minutes at room temperature, followed by two washes with distilled water and air drying.

Synaptosomes (25 μg protein in 80 μL) were added to each well and the plate was centrifuged at 3,000 × g for 1 hour at 4°C without brake to ensure synaptosome attachment. Following centrifugation, an additional 100 μL of assay medium was gently added to each well, bringing the final volume to 180 μL per well. The plate was equilibrated in a non-CO₂ incubator at 37°C for 1 hour before analysis. The mitochondrial stress test protocol involved sequential injection of mitochondrial modulators to assess different aspects of respiratory function: 1) oligomycin (1 μM final concentration) to inhibit ATP synthase and measure ATP-linked respiration; 2) carbonyl cyanide-4-(trifluoromethoxy)phenylhydrazone (FCCP, 2 μM final concentration) to uncouple mitochondrial respiration and determine maximal respiratory capacity; and 3) combined rotenone and antimycin A (0.5 μM each, final concentration) to inhibit Complexes I and III respectively and define non-mitochondrial respiration. Each compound was dissolved in assay medium and loaded into the appropriate injection ports of a Seahorse sensor cartridge that was hydrated overnight in XF Calibrant at 37°C in a non-CO₂ incubator. Oxygen consumption rate (OCR) measurements were taken at baseline (3 measurements over 18 minutes) and following each sequential injection (3 measurements over 18 minutes each) with a mix time of 3 minutes and measurement time of 3 minutes per data point. Respiratory parameters were calculated as follows: basal respiration (baseline OCR minus rotenone/antimycin-A OCR), ATP production (baseline OCR minus oligomycin OCR), maximal respiration (FCCP OCR minus rotenone/antimycin-A OCR) and spare respiratory capacity (maximal respiration minus basal respiration. Data were normalized to cell number. Results represent the mean ± SD from at least three independent synaptosome preparations with 8-12 technical replicates per condition, with statistical comparisons performed between WT and SYNJ2^AAA^ synaptosomes for each respiratory parameter.n=20-24 for each group.

### Electron microscopy of sciatic nerve

For ultrastructural analysis of peripheral nerve morphology, sciatic nerves were isolated from WT and SYNJ2^AAA^ mice (six months) following CO₂ asphyxiation and cervical dislocation. Sciatic nerves were carefully dissected from the sciatic notch to the knee under a stereomicroscope, with particular attention to avoid mechanical damage. Nerves were immediately placed in primary fixative solution containing 2.5% glutaraldehyde and 2% paraformaldehyde in 0.1 M phosphate buffer (pH 7.4) and fixed for 2 hours at 4°C. Following primary fixation, nerve segments were cut into 1-2 mm transverse sections and post-fixed with 1% osmium tetroxide in 0.1 M phosphate buffer for 1 hour at 4°C. Tissue was then dehydrated through a graded ethanol series (50%, 70%, 85%, 95%, and 100% ethanol, 15 minutes each) followed by two changes of propylene oxide for 15 minutes each. Specimens were infiltrated with a 1:1 mixture of propylene oxide and Spurr’s resin for 2 hours at room temperature, followed by pure Spurr’s resin overnight at room temperature with gentle rotation. Tissues were embedded in fresh Spurr’s resin and polymerized at 60°C for 48 hours. Semi-thin sections (1 μm) were cut using a Reichert ultramicrotome and stained with toluidine blue to assess tissue morphology and select appropriate areas for ultrathin sectioning. Ultrathin sections (60-80 nm) were cut with a diamond knife, mounted on copper grids, and stained with uranyl acetate (15 minutes) followed by lead citrate (5 minutes). Sections were examined using a transmission electron microscope (JEOL 1400 or equivalent) operating at 80 kV. For quantitative analysis, systematic random sampling was performed with images captured at 5,000× and 15,000× magnification. At least 10 randomly selected fields per animal were analyzed, with measurements taken from a minimum of 5 animals per genotype. Morphometric parameters assessed included: mitochondrial cristae circularity, size, and number. Data were expressed as mean ± SD and statistical comparisons between WT and SYNJ2^AAA^ groups were performed using unpaired t-tests or Mann-Whitney U tests as appropriate. n=150-175 for each group.

### Metabolite extraction and mass spectrometry analysis

Freshly isolated synaptosomes from WT and SYNJ2^AAA^ mice were immediately processed for metabolite extraction. Synaptosome pellets (200–300 μg protein) were collected by centrifugation, washed three times, and lysed in ice-cold methanol:water (80:20) at dry-ice temperature, with vigorous vortexing for 30 sec. The quantity of metabolite fraction analyzed was normalized to protein concentration determined from a parallel sample. Samples were dried and subjected to liquid chromatography and mass spectrometry (LC-MS) analysis that covers over 200 metabolites by J.M Asara’s group at Beth Israel Deaconess Medical Center, as previously described. Processed data were analyzed using MetaboAnalyst 5.0 (www.metaboanalyst.ca). Analyses were performed on four independent synaptosome preparations. Peak intensity data were uploaded as a CSV file with samples in rows and metabolites in columns, including group labels and sample types. Data preprocessing included: missing value imputation using the k-nearest neighbors method (k=5), log transformation and Pareto scaling for normalization, and outlier detection using principal component analysis (PCA). Quality control was assessed by examining QC sample clustering in PCA score plots. Univariate statistical analysis was performed using unpaired t-tests with false discovery rate (FDR) correction using the Benjamini-Hochberg method. Metabolites with FDR-adjusted p-values <0.05 and fold change >1.5 were considered significantly altered. Multivariate analysis included PCA for data overview and partial least squares discriminant analysis (PLS-DA) to identify discriminating metabolites between groups. Model validity was assessed using 7-fold cross-validation and permutation testing (n=1000). Pathway enrichment analysis was conducted using the pathway analysis module in MetaboAnalyst with the Mus musculus (mouse) pathway library from KEGG. Significantly altered metabolites were mapped to biological pathways using hypergeometric tests for over-representation analysis and pathway topology analysis based on relative betweenness centrality. Pathways with FDR-adjusted p-values <0.05 and pathway impact scores >0.1 were considered significantly enriched. Network visualization was performed to identify metabolic pathway interactions and potential biomarker panels. Results were validated by examining individual metabolite profiles and confirming pathway relevance to synaptic function and SYNJ2 biology. n=9 for each group.

### Lactate measurement

Lactate levels in isolated synaptosomes were measured using the Lactate-Glo™ Assay (Promega), following the manufacturer’s protocol for cell lysates with modifications for synaptosomal preparations. Freshly isolated synaptosomes from WT and SYNJ2^AAA^ mice were resuspended in ice-cold PBS to a defined protein concentration, and 25 μL aliquots were transferred to a 96-well plate. Inactivation Solution (0.6N HCl; 12.5 μL) was added to each well and mixed by shaking for 5 min to lyse synaptosomes, inhibit endogenous enzymatic activity, and degrade NAD(P)H. Samples were then neutralized with 12.5μL Neutralization Solution (1M Trizma base) and mixed by shaking for 30–60 sec. Lactate Detection Reagent (50μL), prepared by combining Reductase, Reductase Substrate, Lactate Dehydrogenase, and NAD in Luciferin Detection Solution per the manufacturer’s instructions, was added to each well, and plates were shaken for 30–60 sec to mix. Following a 60 min incubation at room temperature, luminescence was recorded using a plate-reading luminometer. Buffer-only wells served as negative controls, and a lactate standard curve (10mM lactate stock serially diluted in PBS) was included on each plate to calculate absolute lactate concentrations. Lactate values were normalized to synaptosomal protein concentration, determined in parallel by BCA assay. For each experimental condition, synaptosomes were prepared from at least 6 individual mice per genotype, with technical triplicates performed for each biological sample. Data were expressed as mean ± SD and statistical significance was assessed using unpaired t-tests to compare lactate levels between WT and SYNJ2^AAA^ synaptosomes. The assay was performed within 4 hours of synaptosome isolation to ensure metabolic integrity and minimize lactate degradation. n=9 for each group.

### SYTOX cell viability analysis

Cell viability in WT and SYNJ2^AAA^ neurons was assessed using SYTOX Green nucleic acid stain (Thermo Fisher Scientific). Primary cortical neurons and iPSC-derived neurons were plated on poly-L-lysine coated glass coverslips in 24-well culture dishes and maintained as described above. To induce mitochondrial damage and assess cell viability, neurons were treated with carbonyl cyanide m-chlorophenyl hydrazone (CCCP) dissolved in DMSO for 6 hours at 37°C in culture medium. Control neurons received an equivalent volume of DMSO vehicle (0.1% final concentration). After treatment, culture medium was replaced with fresh medium containing 1 μM SYTOX Green nucleic acid stain and cells were incubated for 15 minutes at 37°C to allow uptake by compromised cells. Images were acquired using a fluorescence microscope (TiEclipse) with a 20× objective lens using appropriate filter sets: DAPI filter for Hoechst 33342 (total nuclei, blue fluorescence) and FITC filter for SYTOX Green (dead/compromised cells, green fluorescence). For each well, at least 10 random fields were captured, ensuring a minimum of 500 total cells per condition for statistical analysis. Images were captured using identical exposure settings within each experiment to allow for direct comparison between conditions. Cell viability analysis was performed using ImageJ software with nuclear segmentation based on Hoechst staining. Total cell number was determined by counting all Hoechst-positive nuclei, while dead cells were identified as nuclei positive for both Hoechst and SYTOX Green staining. Cell viability was calculated as the percentage of cells that were SYTOX-negative (viable cells) relative to the total number of Hoechst-positive nuclei. Dead cell percentage was calculated as (SYTOX-positive cells / total Hoechst-positive cells) × 100. For each experimental condition, coverslips were prepared from at least 4 independent neuronal preparations per genotype, with technical duplicates performed for each CCCP concentration. Data were expressed as mean ± SD and analyzed using two-way ANOVA with Tukey’s multiple comparisons test to compare cell viability between WT and SYNJ2^AAA^ neurons across different CCCP concentrations. n=12 for each group.

### NMDA excitotoxicity assay

Primary cortical neurons and iPSC-derived neurons were plated on poly-L-lysine coated glass coverslips in 24-well culture dishes and maintained as described above. Cells were transfected with pEGFP-C1 plasmid using PEI at a 3:1 PEI:DNA ratio to label viable neurons with green fluorescent protein. At 24 hours post-transfection, baseline GFP-positive cell counts were obtained by capturing fluorescence images from 10 random fields per coverslip using a fluorescence microscope with a 20× objective and FITC filter set. Images were taken using identical exposure settings to ensure consistent quantification across all experimental conditions. For NMDA treatment, culture medium was replaced with pre-warmed Neurobasal medium containing NMDA (100 μM, Tocris Bioscience) plus glycine (10 μM) as a co-agonist to fully activate NMDA receptors. Control neurons received medium containing only glycine (10 μM) without NMDA. Neurons were exposed to NMDA for exactly 10 minutes at 37°C. The medium was rapidly aspirated and neurons were washed three times with warm, Neurobasal medium to completely remove NMDA. Fresh complete medium (Neurobasal with B27, GlutaMAX, and antibiotics) was added, and neurons were returned to the incubator for a 24-hour recovery period to allow excitotoxic cell death to occur. GFP-positive cells were then counted in the same fields that were imaged at baseline to assess neuronal survival. Images were acquired using identical microscope settings and exposure parameters as the baseline images. Cell counting was performed using ImageJ software. Only neurons displaying clear GFP fluorescence with intact cell bodies and visible processes were counted as viable. Neuronal survival was calculated as the percentage of GFP-positive cells remaining at 24 hours post-treatment relative to baseline counts. For each experimental condition, coverslips were prepared from at least 5 independent neuronal preparations per genotype, with technical duplicates for each treatment. Data were expressed as mean ± SD and analyzed using two-way ANOVA with Sidak’s multiple comparisons test to compare neuronal survival between WT and SYNJ2^AAA^ neurons under control and NMDA treatment conditions. Additional analysis included calculation of percent cell death for each condition: [(baseline GFP+ cells - 24h GFP+ cells) / baseline GFP+ cells] × 100. Statistical significance was set at p < 0.05. To validate the assay specificity, selected experiments included co-treatment with the NMDA receptor antagonist APV (50 μM) to confirm that cell death was mediated through NMDA receptor activation. n=9 for each group.

### IncuCyte neurite outgrowth analysis

Neurite outgrowth dynamics were assessed using the IncuCyte S3 Live-Cell Analysis System (Sartorius) under metabolic stress conditions. Primary cortical neurons and iPSC-derived neurons were plated on coated 96-well imaging plates (Greiner Bio-One) and maintained in standard culture medium as described above. Plates were placed in the IncuCyte S3 system housed within a humidified incubator maintained at 37°C with 5% CO₂. Phase-contrast images were automatically captured every 2 hours using a 10× objective lens. For each well, images were acquired from 4 pre-defined regions to ensure comprehensive coverage while maintaining consistent imaging positions throughout the experiment. Automatic focus adjustment was enabled to compensate for potential focal drift during the extended imaging period.

Thirty-six hours after initial plating, culture medium was replaced with glucose-free galactose medium to induce metabolic stress and force reliance on oxidative phosphorylation rather than glycolysis. The galactose medium consisted of Neurobasal-A medium (without glucose, Thermo) supplemented with 10 mM galactose, 1 mM sodium pyruvate, 2 mM L-glutamine. Control wells remained in standard glucose-containing medium for comparison. The media change to galactose was performed gently to minimize mechanical disruption of existing neurites, and wells were visually inspected to ensure no detachment of neurons. Neurite outgrowth analysis was performed using the IncuCyte Neurotrack software module, which automatically identifies cell bodies and traces neurite processes based on phase-contrast morphology. Analysis parameters were optimized for each cell type and included: minimum neurite length of 20 μm, neurite width tolerance of 3-15 μm, and cell body cluster size of 50-500 μm². The software generated multiple readouts including total neurite length per image, number of neurites, mean neurite length per neuron, and neurite branching index. Statistical analysis was performed using two-way ANOVA with repeated measures to compare neurite outgrowth between WT and SYNJ2^AAA^ neurons over time, with Sidak’s multiple comparisons test for post-hoc analysis at individual time points. Additional comparisons included analysis of final neurite parameters at 96 hours using unpaired t-tests between genotypes within each media condition (glucose vs galactose). Data were expressed as mean ± SD, with statistical significance set at p < 0.05. This approach allowed assessment of whether SYNJ2^AAA^ mutations affect neuronal adaptation to metabolic stress and the ability to maintain neurite growth when forced to rely on oxidative metabolism rather than glycolytic energy production. n=20-24 for each group.

### Statistical analysis

All statistical analyses were performed in GraphPad Prism version 10. Data are presented as mean ± SD. Comparisons between two groups used unpaired two-tailed Student’s t-tests, or Mann-Whitney U tests where noted. Comparisons across multiple groups or conditions used two-way ANOVA followed by post hoc multiple comparisons tests. Colocalization was quantified using Manders’ colocalization coefficients (M1 and M2), calculated in Fiji/ImageJ. A p-value < 0.05 was considered statistically significant. Sample sizes for each experiment are reported in the corresponding Methods subsections above.

## Acknowledgements

We thank the IDDRC Gene Manipulation Core, funded by NIH P50 HD105351.

---

**Supplement Figure 1: (A)** Sanger sequencing confirming correct incorporation of the AAA mutation at residues V892, Q934, and L936 in genomic DNA isolated from Synj2^AAA^ mouse brain. Sequences are aligned to the reference mouse genome (NCBI). **(B)** Sanger sequencing confirming correct incorporation of the AAA mutation at residues V892, Q934, and L936 in genomic DNA isolated from Synj2^AAA^ iPSC-derived cortical-like neurons. Sequences are aligned to the reference human genome (NCBI). **C)** qPCR analysis of cDNA from WT and Synj^AAA^ mouse brain and iPSC-derived cortical neurons. Primers targeting Synj2 exon 18 were used to measure total Synj2 mRNA, and primers specific to the mitochondrial-targeting sequence of Synj2a were used to measure the mitochondrial-specific isoform. Compared to WT controls, total Synj2 and Synj2a RNA levels were unchanged. Tubulin was used as a loading control. n=9.

## Notes

### Competing Interest Statement

The authors have declared no competing interest.

