## Supplement Figure 1 for "Synaptojanin 2 is required for neuronal mRNA localization, mitochondrial structure and function, and synaptic metabolism"

A

```
Reference          AGGGTCTTCCAGGAAGTGTCCTCTGTCCAAGGCCCGCTGGATGCCACCGTTGTTGTAAAC
Ms SYNJ2AAA        AGGGTCTTCCAGGAAGTGTCCTCTGTCCAAGGCCCGCTGGATGCCACCGCCGTTGTAAAC
                   *************************************************  *********

Reference          CTCCAGTCTCCAACCCTAGAAGAGAAAAATGAATTTCCAGAGGACCTGCGCACAGAACTT
Ms SYNJ2AAA        CTCCAGTCTCCAACCCTAGAAGAGAAAAATGAATTTCCAGAGGACCTGCGCACAGAACTT
                   ************************************************************

Reference          ATGCAGACTTTGGGGAATTATGGGACGATTATTCTAGTCAGGTAATCCCCGCCCTTGAGA
Ms SYNJ2AAA        ATGCAGACTTTGGGGAATTATGGGACGATTATTCTAGTCAGGTAATCCCCGCCCTTGAGA
                   ************************************************************

Reference          CATATGTTCTGGGGTGTCCGAGGGATCCAAGAGAGCTGCTGCATTAGGTTTCTGTATTGA
Ms SYNJ2AAA        CATATGTTCTGGGGTGTCGGAGGGATCCAAGAGAGCTGCTGCATTAGGTTTCTGTATTGA
                   ************************************************************

Reference          CGAGCTAATCTCCTGTAACACAGGATCAACCAAGGGCAGATGCTGGTGACGTTTGCAGAC
Ms SYNJ2AAA        CGAGCTAATCTCCTGTAACACAGGATCAACCAAGGGGCCATGGCCGTGACGTTTGCAGAC
                   ************************************ . *** . ***************

Reference          AGCCACTCGGCTCTCAGTGTGCTGGATGTGGATGGTATGAAGGTAAGTGGTGTTTAGTCA
Ms SYNJ2AAA        AGCCACTCGGCTCTCAGTGTGCTGGATGTGGATGGTATGAAGGTAAGTGGTGTTTAGTCA
                   ************************************************************
```

B

```
Reference          GGGAGAGGGTTTTCCAGGAAGTGTCCTCCTTCCAGGGCCCCCTGGATGCCACTGTTGTAG
Hu SYNJ2AAA        GGGAGAGGGTTTTCCAGGAAGTGTCCTCCTTCCAGGGCCCCCTGGATGCCACTGCCGTAG
                   ******************************************************  ****

Reference          TAAACCTTCAATCACCGACCTTAGAAGAGAAAAACGAGTTTCCAGAGGACCTGCGTACTG
Hu SYNJ2AAA        TAAACCTTCAATCACCGACCTTAGAAGAGAAAAACGAGTTTCCAGAGGACCTGCGTACTG
                   ************************************************************

Reference          AGCTCATGCAGACCTTGGGGAGTTATGGGACAATTGTTCTTGTCAGGTAACTGCTCCCCT
Hu SYNJ2AAA        AGCTCATGCAGACCTTGGGGAGTTATGGGACAATTGTTCTTGTCAGGTAACTGCTCCCCT
                   ************************************************************

Reference          GGCTGATGTGGGTTCCAGGGGATGTCGATGAGGATCCGTGAGAGCTGCCAGGTCGTTCTT
Hu SYNJ2AAA        GGCTGATGTGGGTTCCAGGGGATGTCGATGAGGATCCGTGAGAGCTGCCAGGTCGTTCTT
                   ************************************************************

Reference          GGCATTGACTTGGAGTATCATTTATTCCAGGATCAACCAAGGGCAGATGCTGGTAACTTT
Hu SYNJ2AAA        GGCATTGACTTGGAGTATCATTTATTCCAGGATCAACCAAGGGGCCATGGCCGTAACTTT
                   *******************************************   ***   ********
```

C

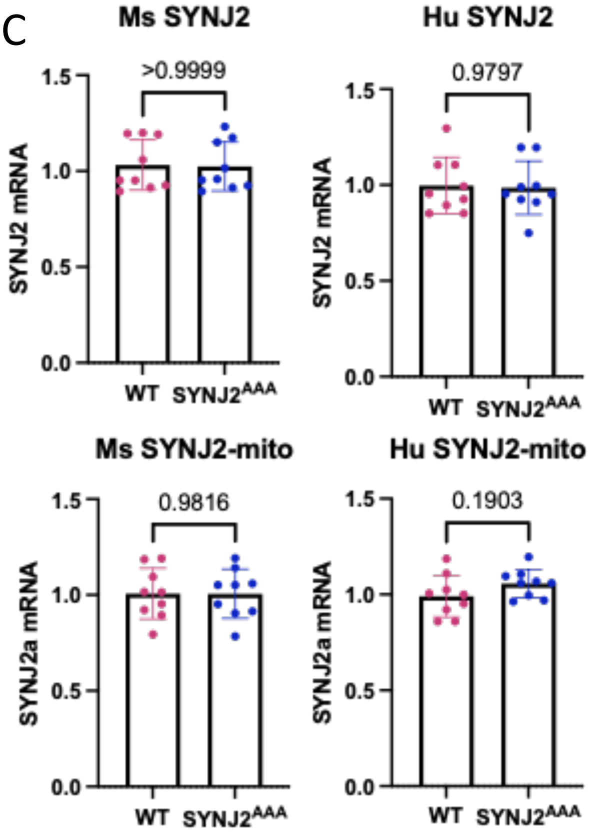
